# MULTI-seq: Scalable sample multiplexing for single-cell RNA sequencing using lipid-tagged indices

**DOI:** 10.1101/387241

**Authors:** Christopher S. McGinnis, David M. Patterson, Juliane Winkler, Marco Y. Hein, Vasudha Srivastava, Daniel N. Conrad, Lyndsay M. Murrow, Jonathan S. Weissman, Zena Werb, Eric D. Chow, Zev J. Gartner

**Affiliations:** University of California San Francisco, Department of Pharmaceutical Chemistry; University of California San Francisco, Department of Anatomy; University of California San Francisco, Department of Cellular and Molecular Pharmacology; Howard Hughes Medical Institute; Chan Zuckerberg BioHub, University of California, San Francisco; Center for Cellular Construction, University of California, San Francisco; Helen Diller Family Comprehensive Cancer Center, San Francisco; University of California San Francisco, Department of Biochemistry and Biophysics; University of California San Francisco, Center for Advanced Technology

## Abstract

We describe MULTI-seq: A rapid, modular, and universal scRNA-seq sample **m**ultiplexing strategy **u**sing **l**ipid-**t**agged **i**ndices. MULTI-seq reagents can barcode any cell type from any species with an accessible plasma membrane. The method is compatible with enzymatic tissue dissociation, and also preserves viability and endogenous gene expression patterns. We leverage these features to multiplex the analysis of multiple solid tissues comprising human and mouse cells isolated from patient-derived xenograft mouse models. We also utilize MULTI-seq’s modular design to perform a 96-plex perturbation experiment with human mammary epithelial cells. MULTI-seq also enables robust doublet identification, which improves data quality and increases scRNA-seq cell throughput by minimizing the negative effects of Poisson loading. We anticipate that the sample throughput and reagent savings enabled by MULTI-seq will expand the purview of scRNA-seq and democratize the application of these technologies within the scientific community.

## INTRODUCTION

Single-cell RNA sequencing (scRNA-seq) has emerged as a powerful technology for probing the heterogeneous transcriptional profiles of multicellular systems. Early scRNA-seq workflows utilized FACS or integrated microfluidics circuits to isolate individual cells and were thus limited to quantifying 10s-100s of single-cell transcriptomes at a time (Tang et al., 2009; Ramsköld et al., 2012; Hashimony et al., 2012). Today, the advent and commercialization of microwell (Gierahn et al., 2017), split-pool barcoding (Rosenberg et al., 2018), and droplet-microfluidics (Macosko et al., 2015; Klein et al., 2015; Zheng et al., 2017) methods has enabled the routine transcriptional analysis of 10^3^-10^5^ cells in parallel. The essential insight enabling these approaches is identical – pools of transcripts are linked to their cell-of-origin via DNA barcodes introduced during reverse transcription and/or ligation. This enormous increase in cell throughput enabled by these methods has catalyzed efforts to catalog the composition of whole organs (The Tabula Muris Consortium et al., 2018) and even entire organisms (Cao et al., 2017; Han et al., 2018). Indeed, ambitious efforts are now underway to create a cell-type atlas for the human body using the latest scRNA-seq techniques (Regev et al., 2017). However, much as research priorities shifted away from describing DNA sequences to functional genomics following the culmination of the Human Genome Project (Lander et al., 2001; ENCODE Project Consortium, 2012), the single-cell genomics field will soon expand beyond descriptive analyses of cell types to mechanistically characterizing how these diverse cell populations interact through space and time to regulate development, homeostasis, and disease.

In order to utilize single-cell sequencing technologies to reveal mechanistic insights into complex multicellular biology, the enormous throughput of scRNA-seq methods must be redirected towards hypothesis testing. This requires integrating dynamical information, many experimental perturbations, and multiple replicates in order to draw strong conclusions. While existing methods are optimally configured to assay many thousands of cells, library preparation practices and the physical constraints of current commercially-available microfluidic devices (e.g., the Fluidigm C1 and 10X Genomics Single-Cell V2 systems) limit analyses to sets of 8 or fewer conditions in a typical scRNA-seq experiment. Experiments that attempt to compare large numbers of samples across multiple single-cell sequencing runs frequently suffer from batch effects (Stegle et al., 2015; Haghverdi et al., 2018). Furthermore, at current prices, the reagent and sequencing costs associated with analyzing large sample numbers is outside the means of typical research groups. One approach to circumvent these challenges would be to sequence large numbers of cells from diverse samples, but with relatively fewer cells from each sample. Encouragingly, recent studies suggest that scRNA-seq data from relatively few cells are sufficient to reconstruct the composition of complex biological tissues (Bhaduri et al., 2017).

Thus, techniques enabling the parallel processing of large sample numbers spanning diverse genetic backgrounds, experimental conditions, and/or time-points will ameliorate known technical limitations while expanding the purview of single-cell genomics to mechanism-oriented biological questions.

Several new multiplexing methods enable parallel sample processing and, thus, more optimal utilization of scRNA-seq cell throughput. These approaches distinguish samples using pre-existing genetic diversity (Kang et al., 2018), or introduce sample-specific DNA barcodes using either genetic (Dixit et al., 2016; Adamson et al., 2016; Jaitin et al., 2016; Aarts et al., 2017; Guo et al., 2018; Shin et al., 2018) or non-genetic (Stoeckius et al., 2017a; Gehring et al., 2018) delivery mechanisms and achieve sample multiplexing via the co-association of sample and transcript barcodes with cell-specific barcodes. Each of these methods has unique liabilities, including sensitivity to proteolytic enzymes necessary to prepare single-cell suspensions, the necessity of reliable surface epitopes for barcoding, compatibility with the harsh transfection or reaction conditions needed to introduce barcodes, poor scalability, or the potential to introduce undesirable secondary perturbations to experiments. Thus, a more generalizable sample barcoding strategy would enable barcodes to be associated with experimental conditions quickly, with high signal-to-noise, and simultaneously on diverse cell lines and tissues from distinct species. This strategy would also be non-perturbative in nature – i.e., to maintain cell viability and endogenous gene expression patterns – and be easily scaled to hundreds or thousands of different samples,

Towards such a generalizable strategy, we report the development of a highly scalable and universal platform for scRNA-seq sample **m**ultiplexing **u**sing **l**ipid-**t**agged **i**ndices (MULTI-seq). MULTI-seq utilizes lipid-modified oligonucleotides (LMOs), which we previously demonstrated to rapidly and stably incorporate into the plasma membrane of live cells via step-wise assembly (Weber et al., 2014). Since LMOs target the plasma membrane, they can be used to barcode any cell or sub-cellular structure with an accessible plasma membrane regardless of species or genetic background. MULTI-seq is non-perturbative, rapid, and involves minimal sample processing, which enables its application to dissociated solid tissues and precious samples. MULTI-seq is also modular in design and, thus, scalable to large sample numbers, as inexpensive and commercially-available unmodified barcode oligonucleotides are localized to membranes via the universal LMO scaffold. We first describe the application of MULTI-seq to multiplex distinct cell lines and culture conditions on a single 10X Genomics Single Cell V2 lane. We then dissociate, barcode, and pool frozen organs from multiple patient-derived xenograft (PDX) mouse models consisting of both mouse and human cells. Finally, we demonstrate scalability by applying MULTI-seq to a 96-sample experimental design where heterogeneous populations of human mammary epithelial cells (HMECs) are stimulated with a panel of growth factors and co-culture conditions.

## RESULTS

### MULTI-seq enables scRNA-seq sample demultiplexing

MULTI-seq localizes sample barcode oligonucleotides to cellular plasma or nuclear membranes via hybridization to a complementary ‘anchor’ LMO targeted to the plasma membrane by a 5’ lignoceric acid amide. Sample barcodes include a 3’ poly-A capture sequence, an 8bp sample barcode, and a 5’ PCR handle necessary for library preparation and anchor LMO hybridization. The off-rate of the anchor LMO from the cell membrane is reduced by subsequent hybridization to an additional ‘co-anchor’ LMO incorporating a 3’ palmitic acid amide that increases the overall hydrophobicity of the complex (Fig. 1B). The same basic strategy can be applied to commercially-available cholesterol-oligonucleotide conjugates, albeit with decreased membrane residence time (Fig. S1D). During droplet microfluidics-based scRNA-seq, cells carry membrane-associated MULTI-seq barcodes into emulsion droplets where, after lysis, the 3’ poly-A domain mimics endogenous transcripts by hybridizing to the oligo-dT regions on co-encapsulated mRNA capture beads. Endogenous transcripts and MULTI-seq barcodes are then linked to a common cell-specific barcode during reverse transcription, which enables sample demultiplexing in the final dataset.

**Figure 1:**
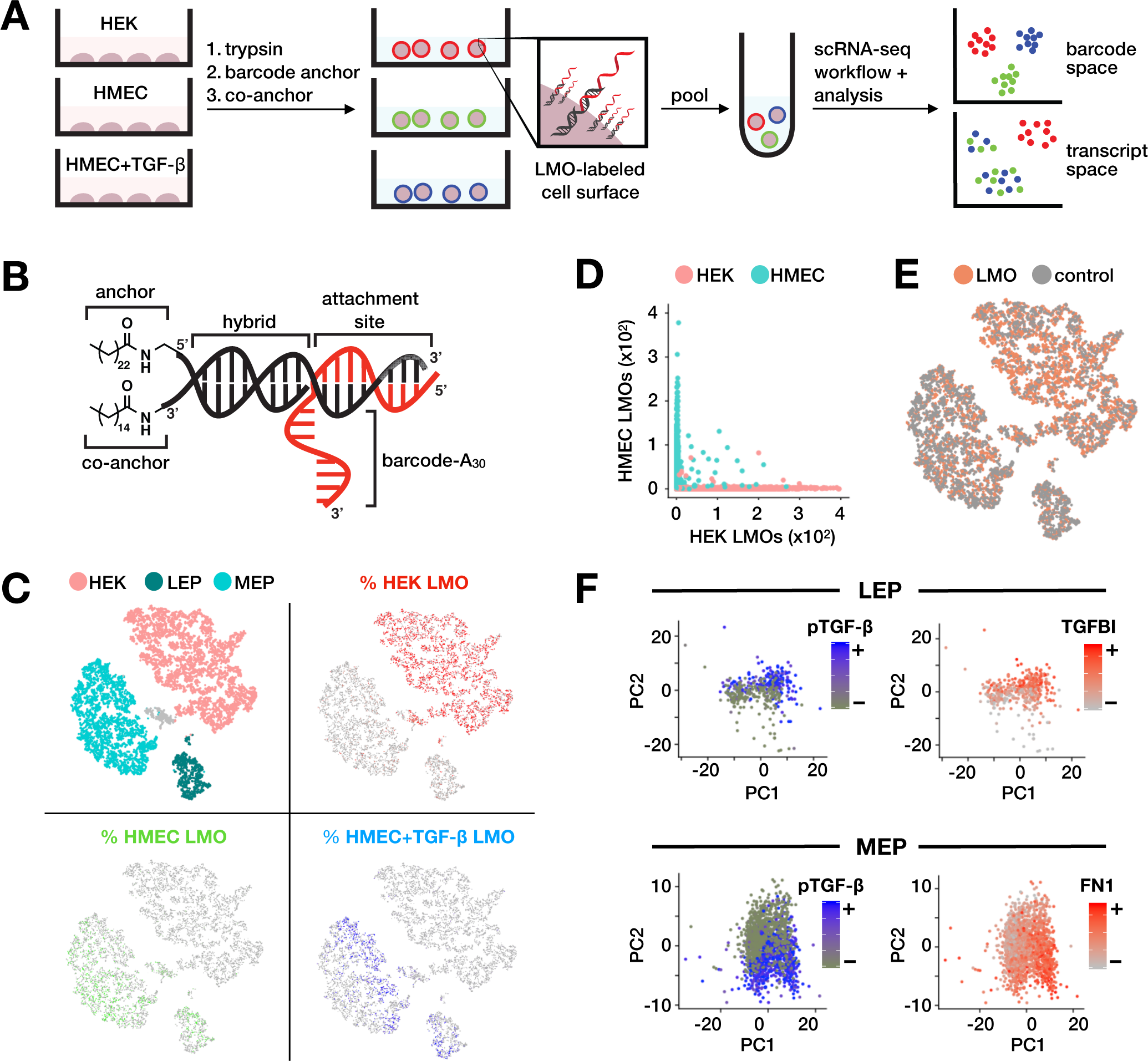
MULTI-seq non-perturbatively demultiplexes cell types and culture conditions. (A) Schematic overview of proof-of-principle MULTI-seq experiment. Three samples (HEKs and HMECs with and without TGF-β stimulation) were barcoded and sequenced alongside unlabeled controls. Labeling involves stepwise assembly of the LMO scaffold on the plasma membrane, where the barcode-hybridized anchor and co-anchor LMOs are added sequentially. Cells are pooled together prior to an augmented scRNA-seq workflow and analysis, producing UMI count matrices corresponding to both gene expression and barcode abundance data. (B) Schematic diagram of the anchor/co-anchor LMO scaffold (black) with hybridized sample barcode oligonucleotide (red). (C) Cell type assignments from marker analysis (top left) largely agree with expected MULTI-seq classification results. HEK-associated clusters are highly enriched for HEK barcodes (top right), while LEP and MEP clusters exhibit enrichment for unstimulated (bottom left) and TGF-β-stimulated (bottom right) barcodes. Cells unclassified via marker analysis (top left, grey) show no barcode specificity. (D) Scatter plot describing the number of barcode UMIs in each cell type. Cell types are highly enriched for their expected barcode and exhibit barcode abundance orthogonality. (E) MULTI-seq barcoded cells (orange) and unlabeled controls (grey) are interspersed in gene expression space. (F) PCA distinguishes stimulated and unstimulated subsets of LEPs and MEPs enriched for transcripts known to be induced (e.g., TGFBI and FN1) in response to TGF-β in HMECs.

Before applying MULTI-seq to a full scRNA-seq experiment, we used flow cytometry to demonstrate that LMOs minimally exchange between cells at 4°C (Fig. S1B,C). Similar experiments were performed using freshly purified cell nuclei (Fig. S1D,E), raising the possibility that this method is equally applicable to single-nucleus RNA sequencing (Habib et al., 2017). Next, we performed a proof-of-principle scRNA-seq experiment to assess whether MULTI-seq can demultiplex distinct cell lines and culture conditions in a non-perturbative fashion. We therefore barcoded and sequenced cultures of HEK293 cells (HEKs) or HMECs with and without stimulation with the growth factor TGF-β on one 10X lane (Fig. 1A). We also sequenced un-barcoded replicates in parallel in order distinguish whether MULTI-seq barcoding influences gene expression. Notably, MULTI-seq barcoding takes 10 minutes at 4° and introduces minimal extra washing steps relative to standard scRNA-seq workflows (Experimental Methods).

We identified clusters in gene expression space according to known markers for HEKs as well as the two primary cellular components of HMECs, myoepithelial (MEPs) and luminal epithelial (LEPs) cells (Fig. 1C, top left; Fig. S2A). Projecting barcode proportions onto gene expression space demonstrates that barcodes are restricted to their intended cell type clusters (Fig. 1C). Furthermore, comparison of barcode counts between cell types also shows minimal background barcode signal, corroborating our previous flow cytometry experiments (Fig. 1D). Importantly, expression profiles for barcoded and control cells are highly similar, demonstrating that MULTI-seq does not alter the cell’s transcriptional state (Fig. 1E; Fig. S2B-C; Supplemental Table S1). To assess whether MULTI-seq demultiplexes culture conditions, we performed sub-clustering and marker analysis on MEPs and LEPs (Fig. S3). Consistent with the culture conditions, LEPs and MEPs classified as TGF-β-stimulated (Computational Methods) express the known TGF-β-induced genes TGFBI and FN1, respectively (Fig. 1F; Hocevar et al., 1999). Collectively, these results illustrate that MULTI-seq accurately demultiplexes distinct cell types and culture conditions without perturbing endogenous gene expression patterns.

### MULTI-seq applied to precious, multi-species PDX samples

An ideal scRNA-seq sample multiplexing platform would be able to simultaneously barcode heterogeneous pools of cells from multiple organisms and tissue types. Moreover, such a technique should involve minimal sample preparation to enable its application to primary and precious tissue sources. To demonstrate these features, we applied MULTI-seq to frozen, dissected tissues from PDX mouse models of triple-negative breast cancer (DeRose et al., 2011; Dobrolecki et al., 2016; Zhang et al., 2013) using a workflow optimized relative to our previous proof-of-principle experiment (Experimental Methods; Fig. S4). Specifically, we barcoded seven samples comprising human primary tumor cells and their associated mouse stroma, matched tumor lung metastases and associated mouse lung stroma, as well as lung stroma from a non-PDX mouse (Fig. 2A). After FACS enrichment for live hCD298^+^ and mCD45^+^ cells, we pooled pre-defined proportions of mouse and human cells together before “super-loading” the 10X Genomics Single Cell V2 system (as in Stoeckius et al., 2017a; Kang et al., 2018), targeting 15,000 cells/lane.

**Figure 2:**
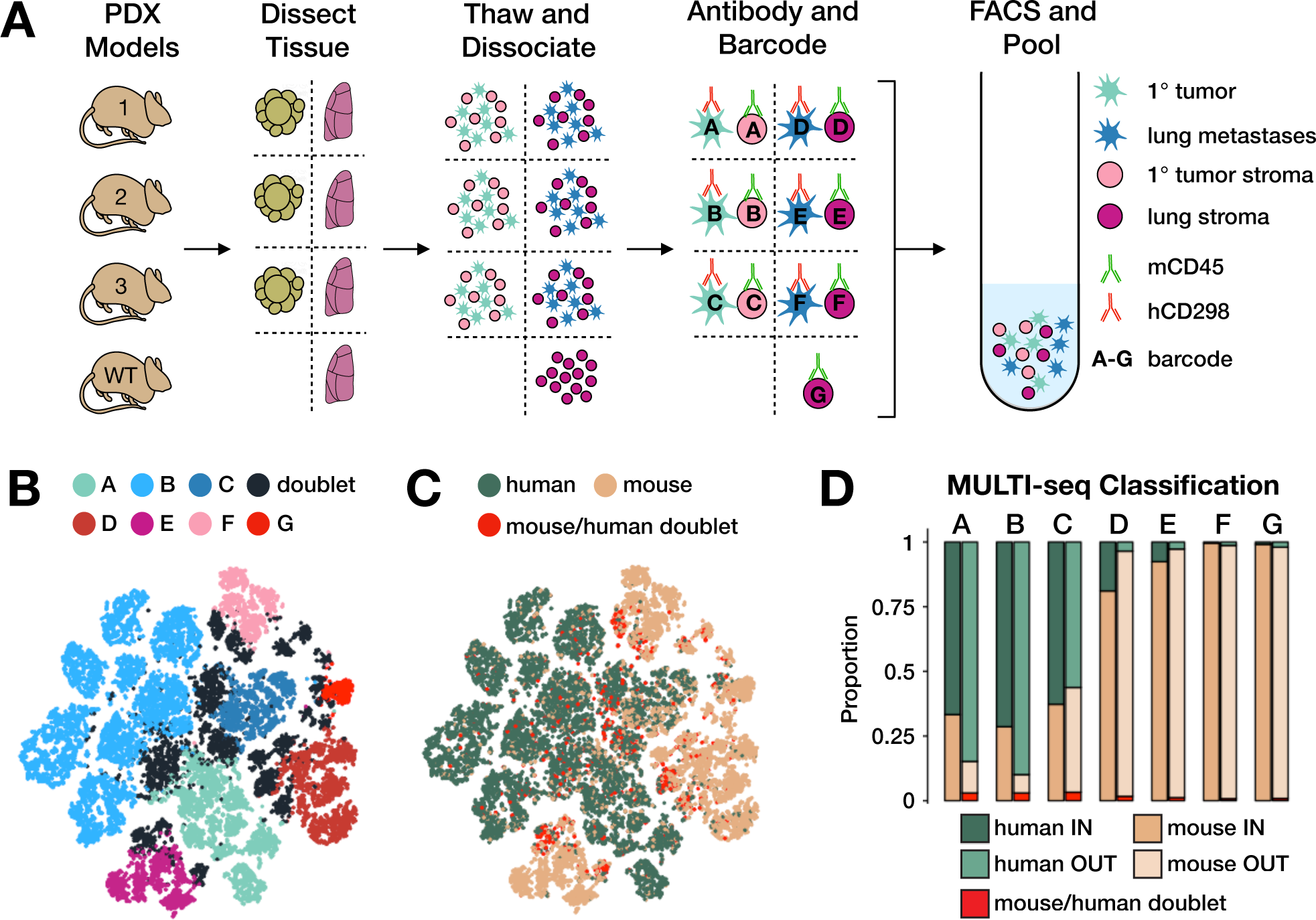
MULTI-seq enables scRNA-seq multiplexing of primary PDX tissue. (A) Schematic overview of PDX experiment. Primary tumors and lung tissue from PDX mouse models were dissected and cryopreserved until the day of the experiment. These tissues were then thawed and dissociated prior to labeling with viability dyes, species-specific antibodies, and sample-specific MULTI-seq barcodes. Live hCD298+ and mCD45+ cells were then FACS-enriched and pooled prior to sequencing. (B) MULTI-seq sample classifications mapped onto barcode space. (C) Species and mouse/human doublet classifications from transcriptome data mapped onto barcode space. (D) Bar plots describing the proportion of mouse (tan) and human (green) cells loaded into the droplet microfluidic device (IN) compared to the species proportions in the final dataset (OUT).

To demultiplex PDX samples in a fashion that both enables doublet identification and takes into account inter-sample barcode variability, we implemented a sample classification workflow inspired by previous work (Stoeckius et al., 2017a; Adamson et al., 2016; Dixit et al., 2016; Computational Methods; Fig. S5). Briefly, we first modeled the probability density function for each sample barcode and identified local maxima corresponding to positive and background cells (As in Adamson et al., 2016; Dixit et al., 2016). Barcode-specific thresholds were then defined by finding the distance between maxima that generates the largest number of singlet classifications across all barcodes. Using this set of barcode-specific thresholds, cells were assigned to a sample group if they surpassed its unique threshold, and cells surpassing more than one threshold were defined as doublets (As in Stoeckius et al., 2017a). Sample demultiplexing illustrates that cells from each MULTI-seq reaction representing both human and mouse cells were detected in the final dataset (Fig. 2B,C). Moreover, comparisons of the proportion of human and mouse cells loaded into the 10X microfluidics device relative to the species proportions in the final dataset generally match expectations (Fig. 2D; Supplemental Table S2). Collectively, these results demonstrate that MULTI-seq can be applied to frozen and solid primary tissue samples while preserving viability and avoiding bias towards specific cell types or species.

### 96-Sample MULTI-seq enables HMEC sample demultiplexing and doublet identification

After demonstrating that MULTI-seq can multiplex scRNA-seq experiments involving both cell lines and primary samples, we next sought to demonstrate the method’s scalability by applying it to 96-distinct samples. To this end, we exposed duplicate cultures consisting of MEPs, LEPs, and a mixture of MEPs and LEPs grown in full M87A media but without EGF (Garbe et al., 2009) to 15 distinct growth factors or growth factor combinations with one control (Fig. 3A). We supplemented this media with growth factors that act within the *in vivo* mammary epithelial microenvironment (e.g., EGF, IGF-1, RANKL, AREG, and WNT4; Brisken, 2013). We barcoded each sample before pooling and “super-loading” four 10X lanes. Pooling resulted in a 24-fold reduction in reagent use relative to standard practices (Experimental Methods), and also minimizes technical noise due to variation between 10X lanes while ensuring that all samples are accounted for in the case of chip failure (e.g., clogged channels, polydisperse droplets, etc.)

**Figure 3:**
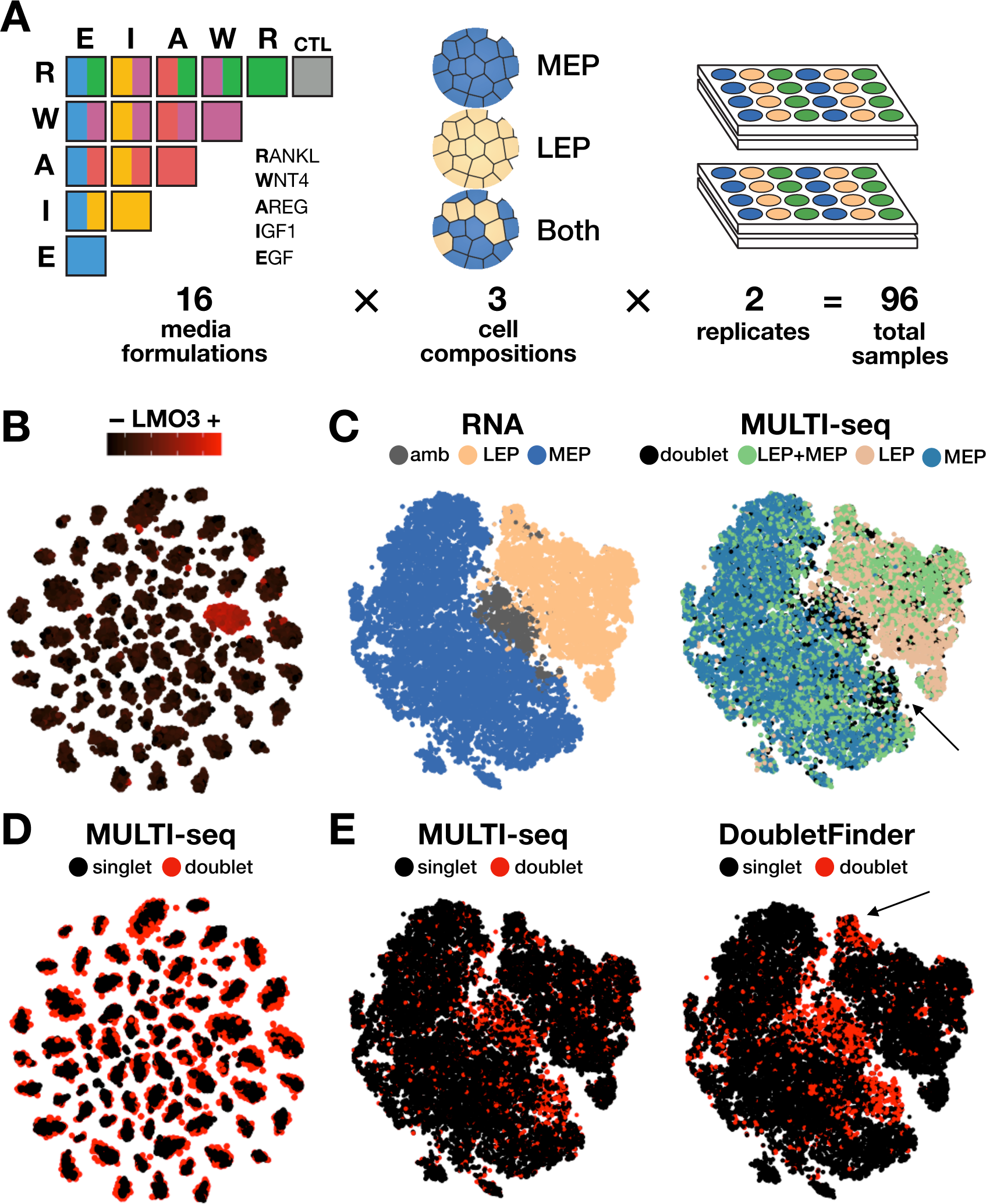
Large-scale MULTI-seq barcoding demultiplexes HMEC culture conditions and identifies doublets. (A) Schematic overview of 96-sample HMEC experiment. 96 distinct HMEC cultures consisting of LEPs alone, MEPs alone, or both cell types together were grown in media supplemented with 15 distinct growth factors or growth factor combinations with one control. (B) Barcode UMI abundance mapped onto barcode space demonstrates that cells cluster according to barcode profiles. LMO barcode #3 is employed as a representative example. (C) Marker analysis identifies LEPs, MEPs, and ambiguous cells in gene expression space (left). MULTI-seq cell-composition classifications (right) match expectations from marker analysis. Region of discordance indicated with the arrow. (D) MULTI-seq doublet classifications mapped onto barcode space illustrates how doublets localize to the peripheries of barcode groups in large-scale sample multiplexing experiments. (E) Doublet classifications produced using MULTI-seq (left) and DoubletFinder (right) mapped onto gene expression space. Region of discordance indicated with the arrow.

After applying our sample classification workflow to this new dataset, we identified 78 high-confidence barcode thresholds which, due to the inclusion of replicates, spanned every distinct experimental condition (Fig. S6A). Each barcode group was associated with an average of 270 cells and each group was enriched for a single barcode ~190-fold above the most abundant off-target barcode and ~1300-fold over the average of all off-target barcodes (Fig. 3B; Fig S6B,C). To test the accuracy of MULTI-seq demultiplexing, we first analyzed the distribution of barcodes associated with different cell compositions (e.g., MEP-alone, LEP-alone, and LEP+MEP samples) in gene expression space. Unsupervised clustering and marker analysis of transcriptome data distinguishes LEPs from MEPs along with a subset of putative doublets expressing markers for both cell types (Fig. 3C, left). MULTI-seq sample classifications match their expected cell type clusters (Fig. 3C, right), while cells co-expressing MEP and LEP markers are predominantly defined as doublets via significant enrichment for multiple MULTI-seq barcodes. Moreover, MULTI-seq doublet classifications are enriched in regions that would have normally been overlooked when predicting doublets using marker genes (Fig. 3C, right). This exemplifies the utility of MULTI-seq and sample multiplexing methods in general for identifying doublets in biological systems where such marker genes are unknown or unavailable.

Encouragingly, MULTI-seq classified 3224 total doublets, which closely matches the expected number of doublets (3046) based on Poisson loading of the 10X microfluidics device. Interestingly, application of an alternative sample classification pipeline (Stoeckius et al., 2017a) to our MULTI-seq data resulted in a 62.4% doublet prediction rate, which is far above the rates estimated by our classification workflow or Poisson statistics (Fig. S6D). We suspect the increased complexity of 96-plex experiments, which alters the relative distribution of singlets and doublets in barcode space compared to smaller-scale experiments (Fig. 3D; Fig. 2B), underlies the requirement for our unique classification pipeline. To further test MULTI-seq doublet classifications, we benchmarked our results against computational identification tools such as DoubletFinder (McGinnis et al., 2018; DePascale et al., 2018; Wolock et al., 2018). DoubletFinder identifies putative doublets by measuring each cell’s proximity to computationally-generated synthetic doublets in gene expression space. DoubletFinder and MULTI-seq doublet predictions significantly overlap in gene expression space, with one putative DoubletFinder false-positive region (Fig. 3E). Collectively, these results indicate that barcode-mediated sample multiplexing is the preferred solution for doublet identification, enabling further increases in cell throughput via droplet-microfluidics device “super-loading.”

### MULTI-seq identifies transcriptional responses to co-culturing and growth factor perturbations

Following sample demultiplexing and doublet removal, we re-analyzed a final scRNA-seq dataset including only MULTI-seq-defined singlets and uncovered three pronounced transcriptional differences driven by variable culture conditions. First, we observed that LEPs co-cultured with MEPs are significantly enriched in the proliferative LEP transcriptional state relative to LEPs cultured alone (Fig. 4A; Supplemental Table S3). In contrast, MEPs were equally proliferative when cultured alone or with LEPs (Fig. 4B). Second, we observed that non-proliferative co-cultured MEPs and LEPs are significantly enriched for TGF-β signaling-induced genes relative to MEPs and LEPs cultured alone (Fig. 4C; Supplemental Table S4). This result indicates that TGF-β signaling in our *in vitro* system cannot be solely maintained via autocrine mechanisms, but, rather, requires paracrine signaling between MEPs and LEPs.

**Figure 4:**
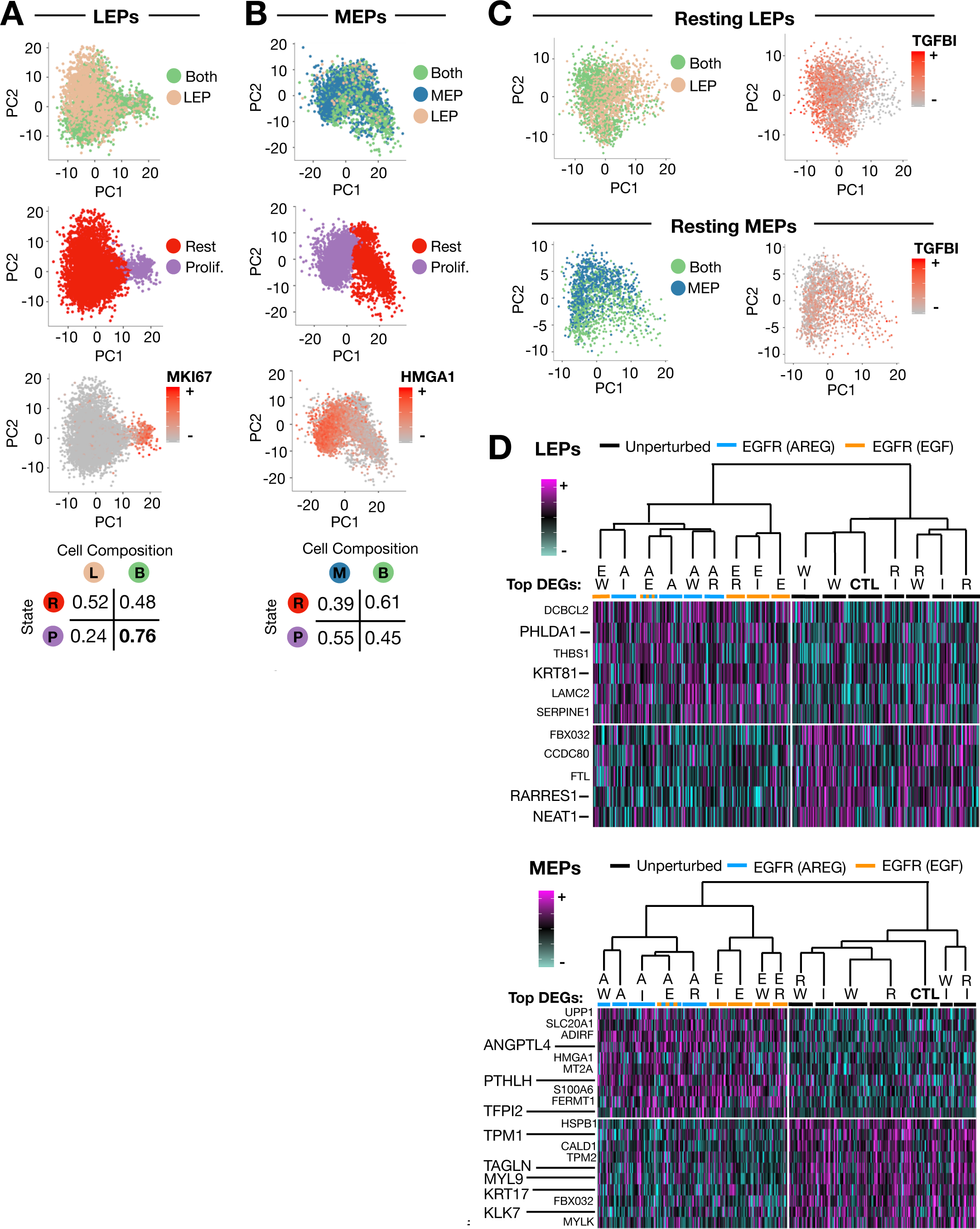
Cell composition and growth factor ulation drive transcriptional changes in HMECs. (A) Gene expression PCA colored by cell composition demonstrates separation of LEPs according to the presence or absence of co-cultured MEPs. LEPs derived from co-cultured samples are highly enriched in a proliferative cluster marked by high levels of MKI67 expression. (B) Gene expression PCA colored by cell composition demonstrates even mixing of MEPs cultured alone or with LEPs across proliferative and resting clusters. Proliferative MEPs are marked by high levels of HMGA1 expression. MEPs assigned to sample barcodes associated with LEP-only culture conditions are due to the inexact nature of EPCAM and CD49f FACS gating (Experimental Methods). (C) Gene expression PCA of subsetted resting LEPs
(top) and MEPs (bottom) colored by cell composition demonstrates enrichment for TGF-α signaling targets in co-culturing conditions. (D) Hierarchical clustering and heat map analysis of LEPs (top) and MEPs (bottom) grouped by growth factor conditions highlights an EGFR signalingrelated transcriptional response present specifically in AREG-and EGF-stimulated HMECs. DEGs = differentially-expressed genes

Third, relative to the co-culture results, we noticed that the transcriptional responses linked to growth factor supplementation were less pronounced. To assess these more nuanced transcriptional effects, we performed hierarchical clustering on the average gene expression profile of MEP and LEP subsets grouped according to growth factor exposure. Interestingly, 100 ng/mL RANKL, WNT4, and IGF-1 did not drive transcriptional signatures that varied significantly from control conditions when added as supplements to M87A (-EGF) growth media (Fig. 4D). In contrast, HMECs exposed to the EGFR ligands AREG and EGF exhibited gene expression profiles that are significantly different from control cells (Supplementary Table S5). Specifically, AREG- and EGF-stimulated MEPs express high levels of the known EGFR-targets (e.g., ANGPTL4, PTHLH, and TFPI2; Savage et al., 2017; Foley et al., 2012; Liao et al., 2015), while unperturbed cells are enriched for known MEP markers involved in contractility (e.g., MYL9, TAGLN, and TPM1/2) and extracellular matrix remodeling (e.g., KRT17 and KLK6/7). AREG- and EGF-stimulated LEPs express high levels of genes known to participate in EGFR signaling negative feedback (e.g., DUSP4; Chitale et al., 2009) or genes up-regulated in HER2^+^ breast cancers (e.g., KRT81 and PHLDA1; Fearon et al., 2018; von der Heyde et al., 2015), while unstimulated LEPs are enriched for known LEP markers (e.g., RARRES1 and NEAT1; Pellacani et al., 2016; Standaert et al., 2014). Collectively, these results demonstrate how MULTI-seq can be applied to study transcriptional responses to varying culture conditions across large numbers of samples.

## DISCUSSION

Recent advances in scRNA-seq cell throughput have facilitated ambitious efforts to catalog the cellular diversity found in whole tissues, organs, and organisms. However, limited sample throughput, high reagent costs, and technical artifacts have slowed the application of scRNA-seq to address more mechanistic biological questions. scRNA-seq sample multiplexing approaches increase the technical and economic feasibility of tackling these questions while removing the confounding influences of batch effects and doublets. We describe here a sample multiplexing strategy – MULTI-seq – that utilizes LMOs to stably localize barcodes to cellular plasma and nuclear membranes.

MULTI-seq has four key characteristics that make it an ideal scRNA-seq multiplexing strategy. First, MULTI-seq sample preparation is rapid, requiring less than 10 minutes to barcode large cell pools at 4°C. This feature, combined with its modular design and the ability to deliver LMOs during proteolytic dissociation, makes MULTI-seq highly scalable and prospectively amenable to automated liquid-handling integration. Further increases in MULTI-seq sample throughput will enable the analysis of drug libraries and/or chemical-genetic screens at single-cell resolution. Unlike traditional small molecule screens that focus on granular read-outs such as cell death or growth rate, highly multiplexed scRNA-seq will provide insight into how small molecules perturb distinct cell types within a multicellular system and drive emergent, population-level responses.

Second, our comparison of barcoded HEKs and HMECs to un-barcoded controls demonstrates that MULTI-seq operates in a non-perturbative fashion on live cells, removing the possibility of incorporating confounding effects associated with fixation, poor viability, genetically-distinct samples, or viral infection. Third, MULTI-seq is universally applicable to all cells with accessible plasma membranes, allowing the same reagents to be applied to multiple cell types from diverse organisms without significant optimization. Together, these two features facilitated our processing of primary dissected tissue from PDX mouse models comprising heterogeneous mouse and human cells that can be challenging to study due to low viability.

Fourth, MULTI-seq exhibits tremendous signal over background (e.g., ~190-fold enrichment for on-target over the most prevalent off-target barcodes), enabling high-confidence sample classification and doublet identification. The ability to detect doublets allows for droplet-microfluidics devices to be “super-loaded”, and thereby further increases scRNA-seq cell throughput by nearly an order of magnitude. Moreover, by benchmarking MULTI-seq doublet classifications against computational doublet identification algorithms, we illustrate how doublets can optimally be handled in scRNA-seq data. Specifically, since many computational doublet prediction algorithms utilize synthetic doublets generated from existing data, false-positives can result when these techniques are applied to datasets with limited transcriptomic diversity (e.g., low numbers of cell types) or cells with gene expression profiles that mimic synthetic doublets (e.g., differentiation intermediates). Such algorithms are also sensitive to false-negatives present in barcode-mediated doublet classifications that arise due to doublets formed from cells labeled with the same sample barcode. Therefore, doublet detection should ideally involve a synergy of computational and molecular approaches, especially in experimental contexts with small numbers of distinct sample barcodes.

In addition to these four desirable technological characteristics, our ability to multiplex a 96-sample HMEC perturbation experiment highlights noteworthy aspects of multiplexed scRNA-seq experimental design. For example, comparison of the transcriptional responses linked to MEP and LEP co-culturing relative to growth factor supplementation demonstrates that transcriptional variation may be dominated by the cell type composition of experimental systems. For instance, co-cultured MEPs and LEPs engage in paracrine-mediated TGF-β signaling that is completely absent in the associated monocultures. In contrast, MEPs and LEPs did not exhibit significant transcriptional changes in rich media supplemented with RANKL, WNT4, and IGF-1 despite the established role of these factors in mammary gland biology. We speculate that the difference in the magnitude of response between co-culturing and small-molecule perturbations can be linked to two distinct phenomena. First, relative to single or combination growth factor perturbations, co-culturing represents a highly complex milieu of stimuli. For example, the pro-proliferative effect of MEP co-culturing in LEPs may be a collective consequence of direct physical interactions and the secretion of extracellular matrix and/or paracrine signaling proteins. Second, rich media formulations likely buffer cells against responding to certain environmental perturbations that the cells are otherwise responsive to *in vivo*. This notion is supported by the observation that the only growth factor supplements that caused significant transcriptional divergence from control cells grown in rich media without EGFR ligands were the EGFR ligands, AREG and EGF. For these reasons, future large-scale scRNA-seq analyses aiming to understand environmental perturbations in *in vitro* systems should be performed in minimal media with careful control of the purity and relative proportions of cell types.

## EXPERIMENTAL METHODS

### Design and synthesis of LMOs and barcodes

Anchor and co-anchor LMO designs were adapted from (Weber et al., 2014). Briefly, the Anchor LMO has a 5’ lignoceric acid modification with two 20-nucleotide domains. The 5’ end is complimentary to the Co-Anchor LMO, which bears a 3’ palmitic acid, and the 3’ end is complimentary to the PCR handle of the Barcode strand. The Barcode oligonucleotide was designed to have three components (as in Stoeckius et al., 2017b):

(1) A 5’ PCR handle for barcode amplification and library preparation, (2) An 8 bp barcode with Hamming distance >3 relative to all other utilized barcodes, and (3) A 30bp poly-A tail necessary for hybridization to the oligo-dT region of mRNA capture bead oligonucleotides (Fig. S6).

Anchor LMO: 5’-GTAACGATCCAGCTGTCACTTGGAATTCTCGGGTGCCAAGG-3’ Co-Anchor LMO: 5’-AGTGACAGCTGGATCGTTAC-3’ Barcode Oligo: 5’-CCTTGGCACCCGAGAATTCCA**NNNNNNNN**A30-3’

### Anchor LMO and co-anchor LMO synthesis

Oligonucleotides were synthesized on an Applied Biosystems Expedite 8909 DNA synthesizer, as previously described (Weber et al., 2014). Hexadecanoic (palmitic) acid, tetracosanoid (lignoceric) acid, N,N-diisopropylethylamine (DIPEA), N,N-diisopropylcarbodiimide (DIC), N,N-dimethylformamide (DMF), methylamine, ammonium hydroxide, and piperidine were obtained from Sigma-Aldrich. HPLC grade acetonitrile (CH_3_CN), triethylamine (NEt_3_), acetic acid, and anhydrous dichloromethane (CH_2_Cl_2_) were obtained from Fisher Scientific. 6-(4-Monomethoxytritylamino)hexyl-(2-cyanoethyl)-(N,N-diisopropyl)-phosphoramidite (5’-Amino-Modifier C6 Phopshoramidite), standard phosphoramidites, and DNA synthesis reagents were obtained from Glen Research. Controlled pore glass (CPG) supports (2-Dimethoxytrityloxymethyl-6-fluorenylmethoxycarbonylamino-hexane-1-succinoyl)-long chain alkylamino-CPG (3’-Amino-Modifier C7 CPG 1000), 5’- Dimethoxytrityl-N-dimethylformamidine-2’-deoxyGuanosine, 3'-succinoyl-long chain alkylamino-CPG (dmf-dG-CPG 1000), and 5’-Dimethoxytrityl-N-Acetyl-2'-deoxyCytidine, 3’-succinoyl-long chain alkylamino-CPG (Ac-dC-CPG 1000) synthesis columns were obtained from Glen Research. All materials were used as received from manufacturer.

For the anchor LMO, after synthesis of the DNA sequence, the 5’ end was modified with an amine using 5’-Amino-Modifier C6 Phopshoramidite (100 mM) and a custom 15-minute coupling protocol. After synthesis of 5′ amino-modified DNA, the MMT protecting group was removed manually on the synthesizer by priming alternately with deblock and dry CH_3_CN at least three times until yellow color disappears. CPG beads were dried by priming several times with dry Helium gas. For the 3′ FMOC-protected amino-modified CPG, prior to oligonucleotide synthesis, the FMOC group was removed by suspending the CPG in a solution of 20% piperidine in dimethylformamide for 10 minutes at room temperature. The beads were then washed three times each with DMF and CH2Cl2. This procedure was repeated twice more to ensure complete deprotection of the FMOC protecting group prior to coupling to the fatty acid. Residual solvent was removed with reduced pressure on a SpeedVac.

Fatty acid conjugation was performed on solid support by coupling the carboxylic acid moiety of the fatty acid to the 3’ or 5’ free amine—lignoceric acid and palmitic acid for the anchor and co-anchor, respectively. The solid support was transferred to a microcentrifuge tube and resuspended in a solution of anhydrous dichloromethane containing 200 mM fatty acid, 400 mM DIPEA, and 200 mM DIC. The microcentrifuge tubes were sealed with parafilm, crowned with a cap lock, and shaken overnight at room temperature. The beads were then washed 3X with CH_2_Cl_2_, 3X with DMF, and 2X CH_2_Cl_2_. Oligonucleotides were then deprotected and cleaved from solid support by suspending the resin in a 1:1 mixture of ammonium hydroxide and 40% methylamine (AMA) for 15 minutes at 65°C with a cap lock followed by evaporation of AMA with a Speedvac system. Cleaved oligonucleotides were dissolved in 0.7 mL of 0.1 M triethylammonium acetate (TEAA) and filtered through 0.2 μM Ultrafree-MC Centrifugal Filter Units (Millipore) to remove any residual CPG support prior to HPLC purification.

Fatty acid modified oligonucleotides were purified from unmodified oligonucleotides by reversed-phase high-performance liquid chromatography (HPLC) using an Agilent 1200 Series HPLC System outfitted with a C8 column (Hypersil Gold, Thermo Scientific) and equipped with a diode array detector (DAD) monitoring at 230 and 260 nm. For HPLC purification, Buffer A was 0.1 M TEAA at pH 7 and buffer B was CH_3_CN. running a gradient between 8 and 95% CH_3_CN over 30 minutes. Pure fractions were collected manually and lyophilized. The resulting powder was then resuspended in distilled water and lyophilized again two more times to remove residual TEAA salts prior to use. Purified fatty acid-modified oligonucleotides were resuspended in distilled water and concentrations were determined by measuring their absorbance at 260 nm on a Thermo-Fischer NanoDrop 2000 series.

### Cell Culture

For proof-of-principle experiments, HEK293 cells were cultured at 37°C with 5% CO2 in Dulbecco’s Modified Eagle’s Medium, High Glucose (DME H-21) containing 4.5 g/L glucose, 0.584 g/L L-glutamine, 3.7 g/L NaHCO_3_, supplemented with 10% fetal bovine serum and 100 μg/mL penicillin/streptomycin. Human mammary epithelial cells (HMECs) were cultured at 37°C with 5% CO_2_ in M87A media (Garbe et al., 2009) with or without 24 hours of stimulation with 5 ng/mL human recombinant TGF-β (Peprotech).

For the 96-sample HMEC experiment, fourth passage HMECs were lifted using 0.05% trypsin+EDTA for 5 minutes. The cell suspension was passed through a 0.45 μm cell strainer to remove any clumps. The cells were washed with M87A media once and resuspended at 10^7^ cells/mL. The cells were incubated with 1:50 APC/Cy-7 anti-human/mouse CD49f (Biolegend, #313628) and 1:200 FITC anti-human CD326 (EpCAM) (Biolegend, #324204) antibodies for 30 minutes on ice. The cells were washed once with PBS and resuspended in PBS with 2% BSA with DAPI at 2-4 million cells/mL. Cells were sorted on BD FACSAria III. DAPI+ cells were discarded. LEPs were gated as EpCAM^hi^/CD49f^lo^ and MEPs were gated as EpCAM^lo^/CD49f^hi^ (Lim et al., 2009; Fig. S7). Notably, this gating strategy results in trace numbers of MEPs and LEPs sorted incorrectly. HMEC sub-populations were sorted into 24-well plates such that wells contained LEPs only, MEPs only, or a 2:1 ratio of LEPs to MEPs. Sorted cell populations were cultured for 48 hours in M87A media before culturing for 72 hours in M87A media (-EGF) supplemented with different growth factors or growth factor combinations. Specifically, M87A media (-EGF) was supplemented with 100 ng/mL RANKL (Peprotech), 100 ng/mL WNT4 (Peprotech), 100 ng/mL IGF-1 (Peprotech), 113 ng/mL AREG (Peprotech), and/or 5 ng/mL EGF (Peprotech) alone or in all possible pairwise combinations.

### Single-cell RNA-seq sample preparation

Distinct sample preparation protocols were employed for the proof-of-principle, 96-plex HMEC, and PDX experiments. For the proof-of-principle experiments, cells were first trypsinized for 5 minutes at 37°C in 0.05% trypsin-EDTA before quenching with appropriate cell culture media. Single-cell suspensions were then pelleted for 4 minutes at 160 x g and washed once with PBS before resuspension in 90 μL of a 200nM solution containing equimolar amounts of anchor LMO and sample barcode oligonucleotides in PBS. Anchor LMO-barcode labeling was performed for 5 minutes on ice before 10 μL of 2μM co-anchor LMO in PBS was added to each cell pool. Following gentle mixing, the labeling reaction was continued on ice for another 5 minutes before cells were washed twice with PBS, resuspended in PBS with 0.04% BSA, filtered and pooled before emulsion using the 10X Genomics Single Cell V2 system.

For the 96-plex HMEC experiment, LMO labeling was performed during trypsinization in order to minimize wash steps and thereby limit cell loss and preserve cell viability. Specifically, HMECs cultured in 24-well plates were labeled for 5 minutes at 37°C and 5% CO_2_ in 190 μL of a 200nM solution containing equimolar amounts of anchor LMO and sample barcode oligonucleotides in 0.05% trypsin-EDTA. 10 μL of 4uM co-anchor LMO in 0.05% trypsin-EDTA was then added to each well and labeling/trypsinization was continued for another 5 minutes at 37°C and 5% CO2 before quenching with appropriate cell culture media. Cells were then transferred to a 96-well plate for washing with 0.04% BSA in PBS. Finally, cells were pooled into a single aliquot, filtered through a 0.45 μm cell strainer, and counted before generating emulsions using the 10X Genomics Single Cell V2 system. The current cost for one 10X microfluidics lane-worth of reagents is ~$1250. For this experiment, we split our pool of 96 samples across 4 10X microfluidics lanes for $5000. In comparison, analyzing 96 samples without multiplexing (i.e., one sample/lane) would therefore cost $120,000.

For the PDX experiment, tissues from primary tumor, lung metastases, and normal lung from PDX models HCI-001, HCI-002 (Derose et al., 2011) and HCI-4272 (Zhang et al., 2013) were generated in NOD-*scid* gamma (NSG) mice as described previously (Lawson et al., 2015). The UCSF Institutional Animal Care and Use Committee (IACUC) reviewed and approved all animal experiments. Frozen tissue was dissociated in digestion media containing 50 μg/mL Liberase TL (Sigma-Aldrich) and 2×10^4^ U/mL DNase I (Sigma-Aldrich) in DMEM/F12 (Gibco) using standard GentleMacs protocols. Single cell suspensions were stained for FACS sorting with Zombie NIR (BioLegend, #423105) and the following antibodies: Fc-block (Tonbo, #70-0161-U500), anti-mouse TER119 (ThermoFisher, #11-5921-82), anti-mouse CD31 (ThermoFisher, #11-0311-85), anti-mouse CD45 (Tonbo, #75-0451-U100), anti-mouse MHC-I (eBioscience, #17-5999-82) and anti-human CD298 (BioLegend, #341704). MULTI-seq labeling was performed using 100uL of a 2.5uM solution containing equimolar amounts of anchor LMO and sample barcode oligonucleotides in PBS. LMO labeling was performed for 5 minutes on ice before 20uL of 15uM co-anchor LMO in PBS was added to each cell pool. LMO labeling was continued for another 5 minutes on ice before cells were washed once with PBS containing 2% FBS prior to live-cell enrichment and separation of mouse CD45^+^ and human tumor cells (CD298^+^ mTER119^−^ mCD31^−^ mMHC-I^−^) via FACS, as described previously (Lawson et al., 2015). MULTI-seq indexed samples were then filtered, counted, and pooled before generating emulsions using the 10X Genomics Single Cell V2 system.

### scRNA-seq Library Preparation

Sequencing libraries were prepared using a custom protocol based on the 10X Genomics Single Cell V2 (10X Genomics, 2017) and CITE-seq (Stoeckius et al., 2017b) workflows. Briefly, the 10X workflow was followed up until cDNA amplification, where 1 μL of 2.5 μM MULTI-Seq additive primer (sequence below) was added to the cDNA amplification master mix. This primer increases barcode sequencing yield by enabling the amplification of barcodes that successfully primed reverse transcription on mRNA capture beads but were not extended via template switching (Fig. S8C). Following amplification, barcode and endogenous cDNA fractions were separated using a 0.6X SPRI size selection. The endogenous cDNA fraction was then processed according to the 10X workflow until sequencing on two HiSeq 4000 lanes (proof-of-principle) or one Nova-Seq lane (96-sample HMEC and PDX).

MULTI-seq Additive Primer: 5’-CCTTGGCACCCGAGAATTCC-3’

Contaminating oligonucleotides remaining from cDNA amplification were then removed from the barcode fraction using an established small RNA enrichment protocol (Beckman Coulter). Specifically, we increased the final SPRI ratio in the barcode fraction to 3.2X reaction volumes and added 1.8X reaction volumes of 100% isopropanol (Sigma-Aldrich). Beads were then washed twice with 400uL of 80% ethanol and allowed to air dry for 2-3 minutes before elution with 50.5μL of Buffer EB (Qiagen, USA). Eluted barcode cDNA was then quantified using QuBit before library preparation PCR (95°C, 5’; 98°C, 15”; 60°C, 30”; 72°C, 30”; 8 cycles; 72°C, 1’; 4°C hold). Each reaction volume was a total of 50μL containing 26.25μL KAPA HiFi master mix (Roche), 2.5μL TruSeq RPIX primer (Illumina), 2.5μL TruSeq Universal Adaptor primer (Illumina), 3.5ng barcode cDNA and nuclease-free water.

### TruSeq RPIX

5’-CAAGCAGAAGACGGCATACGAGAT**NNNNNN**GTGACTGGAGTTCCTTGGCACCCGAGAATTCCA-3’

TruSeq P5 Adaptor:

5’-AATGATACGGCGACCACCGAGATCTACACTCTTTCCCTACACGACGCTCTTCCGATCT-3’

Following library preparation PCR, remaining sequencing primers and contaminating oligonucleotides were removed via a 1.6X SPRI clean-up and sequencing on one HiSeq4000 lane. Representative Bionalayzer traces at different stages of MULTI-seq library preparation are documented in Fig. S8.

### Live-Cell LMO Exchange Experiments

The BD FACSCalibur instrument was used to performed analytical flow cytometry experiments assessing the kinetics of LMO and cholesterol-modified oligonucleotide (CMO) exchange on live HEK293 cells (Fig. S1A-C). Data analysis was performed in FlowJo and R. Identical sample preparation protocols were employed for the proof-of-principle scRNA-seq experiment (discussed above) and live-cell flow cytometry experiments, with one key exception. Instead of pre-hybridizing anchor LMOs or CMOs to barcode oligonucleotides, anchor LMOs or CMOs were pre-hybridized to equimolar concentrations of FAM-or AlexaFluor647-conjugated oligonucleotides. Fluorophore-conjugated oligonucleotides were identical to the barcode oligonucleotide 5’ PCR handle and did not include the barcode or poly-A regions. FAM- and Alexa647-labeled HEK293 cells or nuclei were mixed immediately prior to analysis and kept on ice for 2 hours in PBS with 0.04% BSA.

### Nuclei Isolation and LMO Exchange Experiments

Nuclei were isolated from HEK293 cells using a protocol adapted from 10x Genomics. Briefly, HEK293 cells were cultured, trypsinized, and washed once with PBS. Cells were pelleted (300 rcf, 4˚C, 4 minutes) and suspended in chilled lysis buffer (0.5% Nonidet P40 Substitute, 10 mM Tris-HCl, 10 mM NaCl, and 3 mM MgCl_2_ in milliQ water) to a density of 2.5 × 10^6^ cells/mL. Lysis proceeded for 5 minutes on ice, after which the lysate was pelleted (500 rcf, 4˚C, 4 minutes) and washed three times in chilled resuspension buffer (1X PBS, 2% BSA). Nuclei were then diluted to a concentration of ~10^6^ nuclei/mL prior to LMO labeling, as described above. Following LMO labeling, nuclei were washed times in 1mL resuspension buffer (500 rcf, 4 minutes). LMO exchange experiments were performed as described previously.

## COMPUTATIONAL METHODS

### scRNA-seq Data Processing

Expression library FASTQs were processed using CellRanger (10X Genomics) and aligned either to the hg19 or concatenated hg19-mm10 reference transcriptomes. High-confidence cells were distinguished from background using a nUMI cut-off of 1000. MULTI-seq barcode library FASTQ files were converted into a barcode UMI count matrix using CITE-seq Count (https://github.com/Hoohm/CITE-seq-Count).

### MULTI-seq Sample Classification

For the 96-sample HMEC and PDX experiments, sample classification was performed using a workflow inspired by previous scRNA-seq multiplexing approaches (Stoeckius et al., 2017a; Adamson et al., 2016; Dixit et al., 2016; Fig. S5). First, raw barcode reads were log2-transformed before barcode abundance normalization via mean subtraction. Following normalization, the probability density function (PDF) for each barcode was defined by applying the ‘approxfun’ R function to the Gaussian kernel density estimation produced using the ‘bkde’ function from the ‘KernSmooth’ R package. We then sought to classify cells according to the assumption that groups of cells that are positive and negative for each barcode should manifest as local PDF maxima. To this end, we trimmed the top and bottom 0.1% of data from each barcode set and chose the lowest and highest maxima as initial solutions. To avoid noisy maxima identification, we then adjusted the low maxima to the maxima with the largest number of associated cells.

With these positive and negative approximations in hand, we next sought to define barcode-specific thresholds. To find the best inter-maxima quantile for threshold definition (e.g., an inter-maxima quantile of 0.5 corresponds to the mid-point), we iterated across 0.01 quantile increments and chose the value that maximized the number of singlet classifications. Optimal inter-maxima distances vary across different MULTI-seq datasets and likely reflect technical noise resulting from variable cell numbers and labeling efficiency between samples. Sample classifications were then made using these barcode-specific thresholds by discerning which thresholds each cell surpasses, with doublets being defined as cells surpassing >1 threshold. Negative cells (i.e., cells surpassing 0 thresholds) were discarded. The process then was repeated on the remaining cells, typically for a total of 3 rounds, until no more cells were classified as negatives. Barcode visualizations using t-SNE were generated using the ‘Rtsne’ function with the ‘initial_dims’ argument set to the total number of unique barcodes.

For the proof-of-principle HEK and HMEC experiment, a simpler classification scheme was used. Specifically, raw barcode counts were first converted to proportions before cells were assigned to HEK, stimulated HMEC or unstimulated HMEC samples according to whichever barcode represented >50% of the total barcode UMIs. Such a classification strategy precludes doublet identification and is sensitive to inter-barcode variability. However, for low-sample experiments where doublets are not a large concern, it is appropriate.

### Expression Library Analysis

CellRanger outputs were analyzed using the ‘Seurat’ R package, as described previously (Butler et al., 2018). Stastically-significant PCs were selected using inflection point estimation on corresponding PC elbow plots. Cell types were defined using Louvian clustering with established marker genes. Analysis of transcriptional responses due to variable culture conditions (e.g., cell type compositions and growth factors for the 96-sample HMEC experiment) were performed using PCA to allow for gene loading interpretation. Genes specific to sample assignments were defined using the ‘bimod’ (REF) and ‘roc’ arguments in the ‘FindMarker’ function. Sample groups exhibiting correlated gene expression profiles were defined using the ‘BuildClusterTree’ function.

## DATA AVAILABILITY

Raw gene expression and barcode count matrices were uploaded to the Gene Expression Omnibus (GSE…). An R implementation of the MULTI-seq sample classification pipeline can be found at https://github.com/chris-mcginnis-ucsf/MULTI-seq.

## ACKNOWLEDGMENTS

We thank M.T. Lewis (UCSF) and A. Welm (University of Utah) or providing PDX models developed by their groups. We also thank M. Owyong, A. Abisoye Ogunniyan, C. Diadhiou, J. Garbe, and J. Hu (UCSF) for technical support.

## AUTHOR CONTRIBUTIONS

E.D.C. and Z.J.G. conceptualized the method. C.S.M. and D.M.P. designed experiments, synthesized LMOs, and optimized the method. C.S.M., D.M.P., and D.N.C. performed analytical flow cytometry experiments. C.S.M., D.M.P., and J.W. performed scRNA-seq experiments. Z.W. and J.S.W. provided tissue and computational resources, respectively. V.S. performed FACS for 96-plex HMEC scRNA-seq experiment. C.S.M. and L.M.M. performed bioinformatics analysis. C.S.M., M.Y.H., and J.W. implemented the sample classification pipeline. C.S.M., D.M.P., Z.J.G. and E.D.C. wrote the manuscript.

## DECLARATION OF INTERESTS

Z.J.G, E.D.C., D.M.P., and C.S.M. have filed patent applications related to the MULTI-seq barcoding method. The contents of this manuscript is solely the responsibility of the authors and does not necessarily represent the official views of the National Institutes of Health.

## FUNDING

This research was supported in part by grants from the Department of Defense Breast Cancer Research Program (W81XWH-10-1-1023 and W81XWH-13-1-0221), NIH (U01CA199315 and DP2 HD080351-01), the NSF (MCB-1330864), and the UCSF Center for Cellular Construction (DBI-1548297), an NSF Science and Technology Center. Z.J.G is a Chan-Zuckerberg BioHub Investigator. D.M.P. is supported by the NIGMS of the National Institutes of Health (F32GM128366). L.M.M is a Damon Runyon Fellow supported by the Damon Runyon Cancer Research Foundation (DRG-2239-15). J.W. and M.Y.H. are supported by EMBO long-term post-doctoral fellowships (ALTF-159-2017 and ALTF-1193-2015, respectively).

---

**Figure S1:**
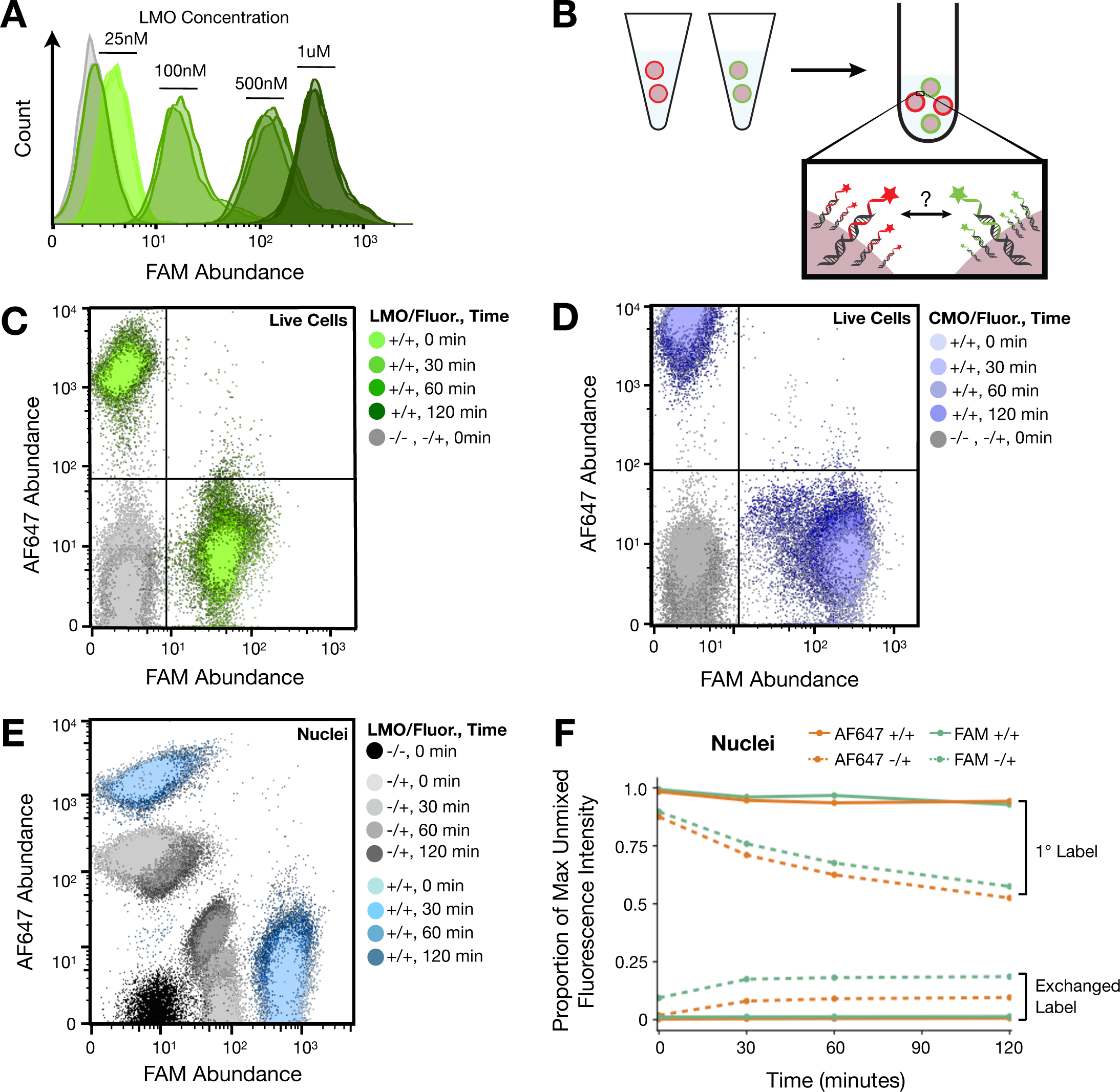
Flow cytometry demonstrates robust LMO labeling efficiency and negligible exchange kinetics at 4˚C on living and nuclei, related to Figure 1. (A) MULTI-seq live-cell labeling efficiency varies predictably across a titration curve of anchor and co-anchor LMO concentrations. (B) Schematic overview of LMO and cholesterol-modified oligonucleotide (CMO) exchange experiments. Cells or nuclei were labeled with LMOs or CMOs hybridized to AF647- or FAM-conjugated oligonucleotides prior to mixing. Mixed populations were kept on ice and analyzed using flow cytometry every 30 minutes for 2 hours. (C) Time-course analysis of LMO exchange following mixing of live cell populations labeled with FAM- or AF647-conjugated barcode oligonucleotides. Control samples receiving nothing (-/-) or fluorophore-conjugated barcode oligonucleotides alone -/+) exhibit minimal background signal relative to samples receiving both LMO and fluorophore (+/+). FAM+ and AF647+ cell populations exchange barcodes at a negligible frequency over 2 hours. (D) Time-course analysis of cholesterol-modified oligonucleotide (CMO) exchange, as depicted in Fig. S1B. Signal loss is more pronounced in CMO-labeled samples than in LMO-labeled samples. (E) Time-course analysis of LMO exchange in nuclei. Control (-/-) samples exhibit no background signal while samples receiving fluorphore alone (-/+) exhibit higher background than live cell experiments. Samples receiving both LMO and fluorophore (+/) are labeled with higher efficiency and exchange less rapidly. (F) Quantification of results in Fig. S1E. Normalization of fluorescence intensity to levels present in unmixed fluorophore-only (-/+) and LMO plus fluorophore +/+ samples illustrates that LMO plus fluorophore samples retain barcodes more robustly over time compared to fluorophore-only controls.

**Figure S2:**
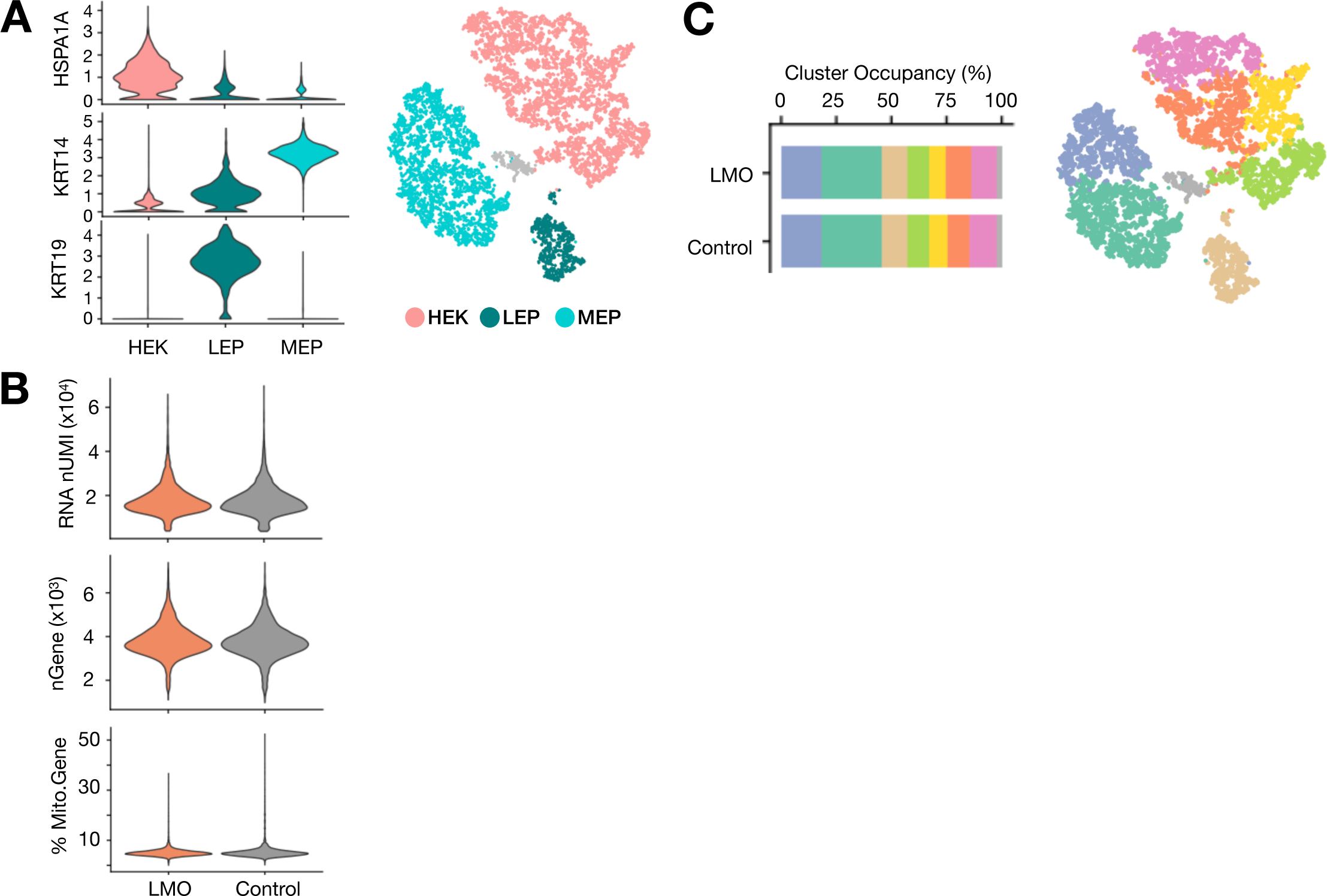
MULTI-seq preserves endogenous gene expression, related to Figure 1. (A) Violin plots describing the expression distribution for marker genes used to delineate HEKs (HSPA1A), MEPs (KRT14), and LEPs (KRT19) as shown in Fig. 1B. (B) Violin plot comparing the number of UMIs, number of detected genes, and percentage of mitochondrial gene expression in MULTI-seq and control cell populations. (C) Unsupervised clustering identifies 8 distinct cell sub-populations. Cluster occupancy rates for equal numbers of randomly sampled barcoded and control HEKs and HMECs illustrates that MULTI-seq does not influence endogenous gene expression.

**Figure S3:**
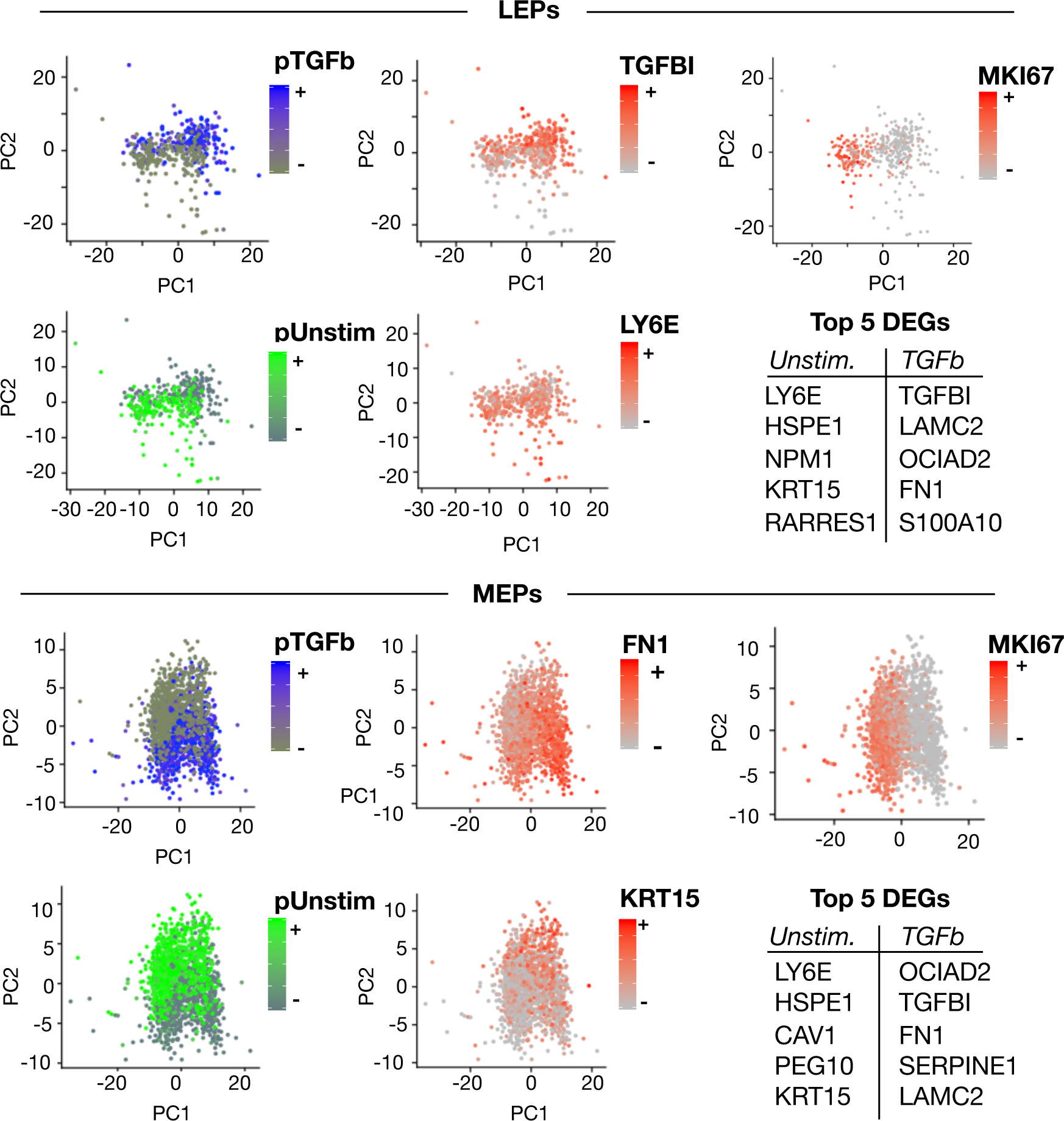
PCA delineates transcriptional differences due to TGF-β-stimulation in subsetted MEPs and LEPs, related to Figure 1. Marker analysis of stimulated and unstimulated MEPs and LEPs uncovers differentially expressed genes between culture conditions that recapitulate known TGF-β targets. Stimulated and unstimulated subsets are resolvable in PC space, and the top five differentially expressed genes for each subset match known TGF-β functions related to microenvironment remodeling (e.g., TGFBI, FN1, LAMC2), as well as acknowledged regulatory interactions (e.g., KRT15, LY6E, SERPINE1). PC1 primarily distinguishes MEPs and LEPs according to proliferation status, as is demonstrates by MKI67 expression enrichment in PC space, whereas PC2 distinguishes TGF-β induction status.

**Figure S4:**
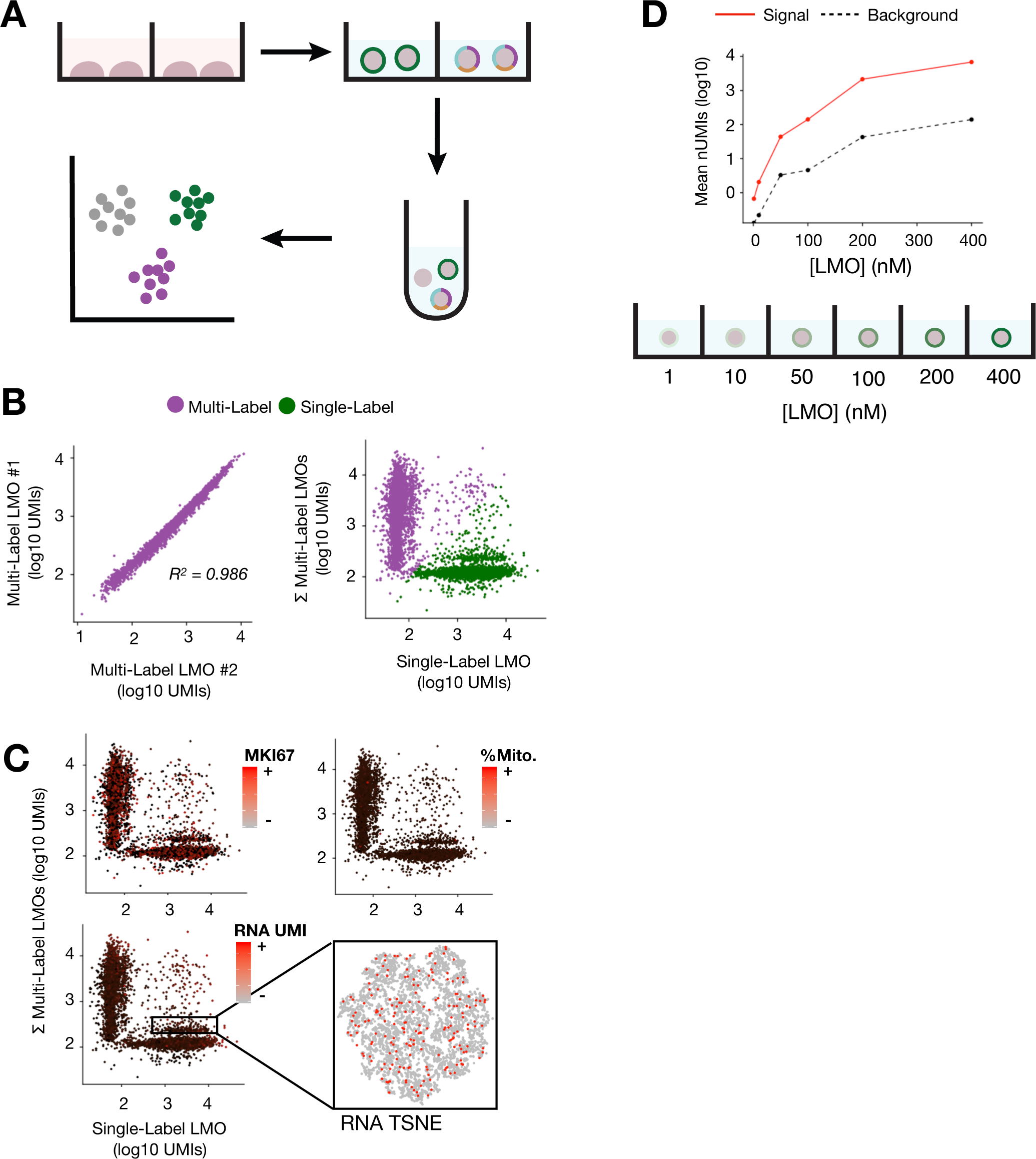
Optimized MULTI-seq workflow enables combinatorial indexing, related to Figure 2. (A) Schematic overview of combinatorial indexing experiment. HEKs were labeled either with an equimolar combination of three barcode LMOs or with a singular barcode LMO. (B) Barcode UMIs in multi-labeled HEKs are highly correlated, suggesting variability in labeling efficiency is primarily biological in nature. Comparison of single-labeled and multi-labeled HEKs demonstrates the orthogonality of labeling. (C) Combinatorial indexing experiment exhibits bimodal background barcode distributions. Exploration of gene expression features that cause bimodality do not yield any clear correlations. Bimodality cannot be linked to changes in cell size due to cell cycle (as measured by MKI67), changes in cell size manifesting as increased RNA content, or apoptotic cells (as measured by the percentage of mitochondrial gene expression). Moreover, the region of relatively high background barcode signal cannot be traced to any particular cell state. (D) MULTI-seq barcode abundances vary predictably across a titration series of anchor and co-anchor LMO concentrations.

**Figure S5:**
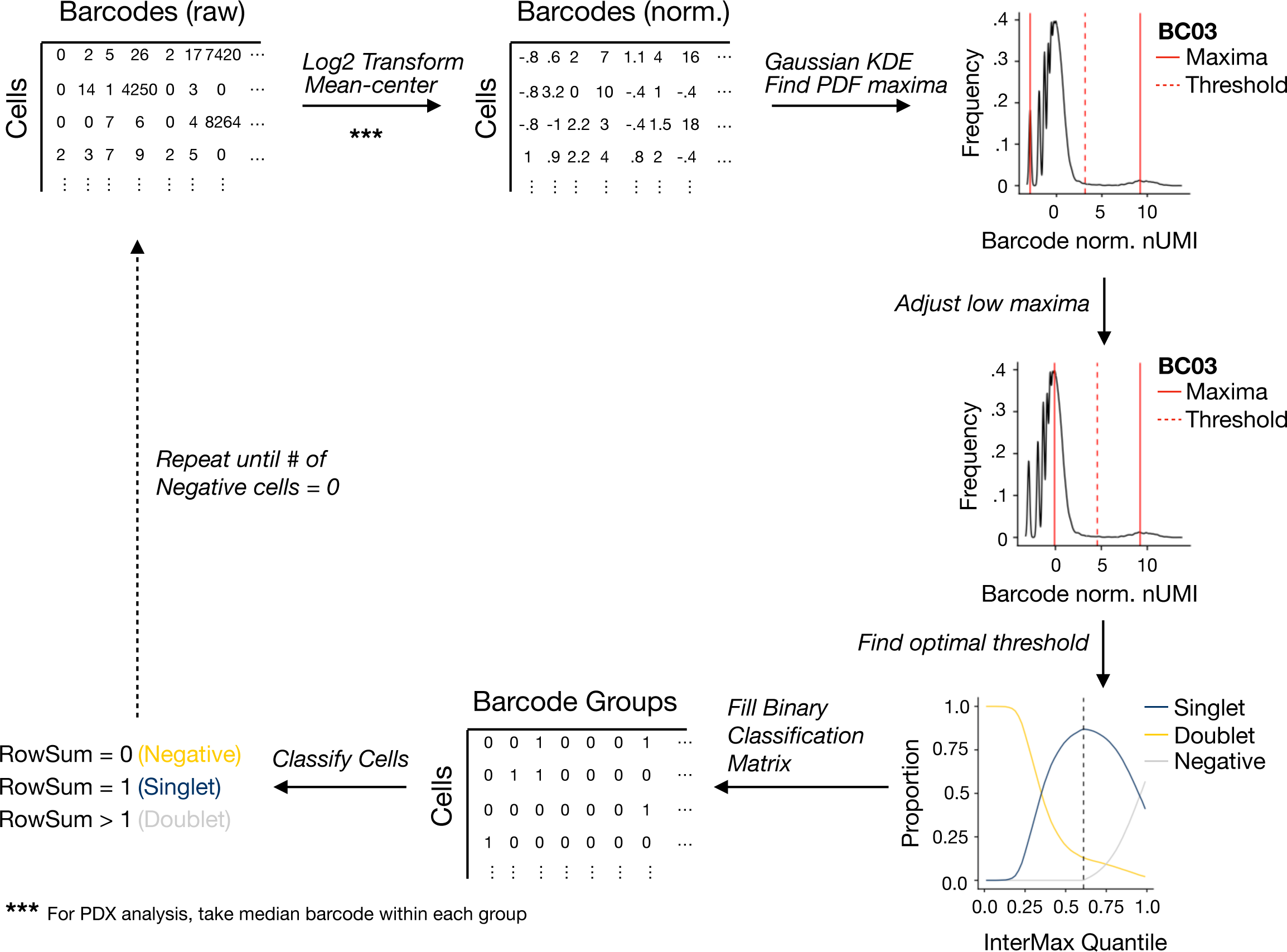
MULTI-seq sample classification workflow, related to Figure 2. Raw barcode UMI count matrices were normalized via Log2 transformation and barcode-oriented mean centering. Using normalized barcode counts, the probability density function (PDF) for each barcode is then defined using Gaussian kernel density estimation (KDE). The lowest and highest local maxima in each PDF are then defined, serving as approximations for cell populations negative or positive for each barcode, respectively. The low maxima is then adjusted to the maxima below the initial threshold with the highest density. Following adjustment, the optimal quantile distance between maxima is determined across all barcodes by finding the quantile which produces the maximum number of singlet classifications. This quantile is then used to set barcode-specific thresholds, which are subsequently utilized to generate a binary classification matrix in which cells are assigned a ‘1’ if they surpass a given threshold. The row-sums of this classification matrix are then used to classify cells, where negative cells, singlets and doublets surpass 0, 1, and >1 threshold, respectively. The pipeline is repeated until all cells are classified as singlets or doublets, with negative cells removed between iterations.

**Figure S6:**
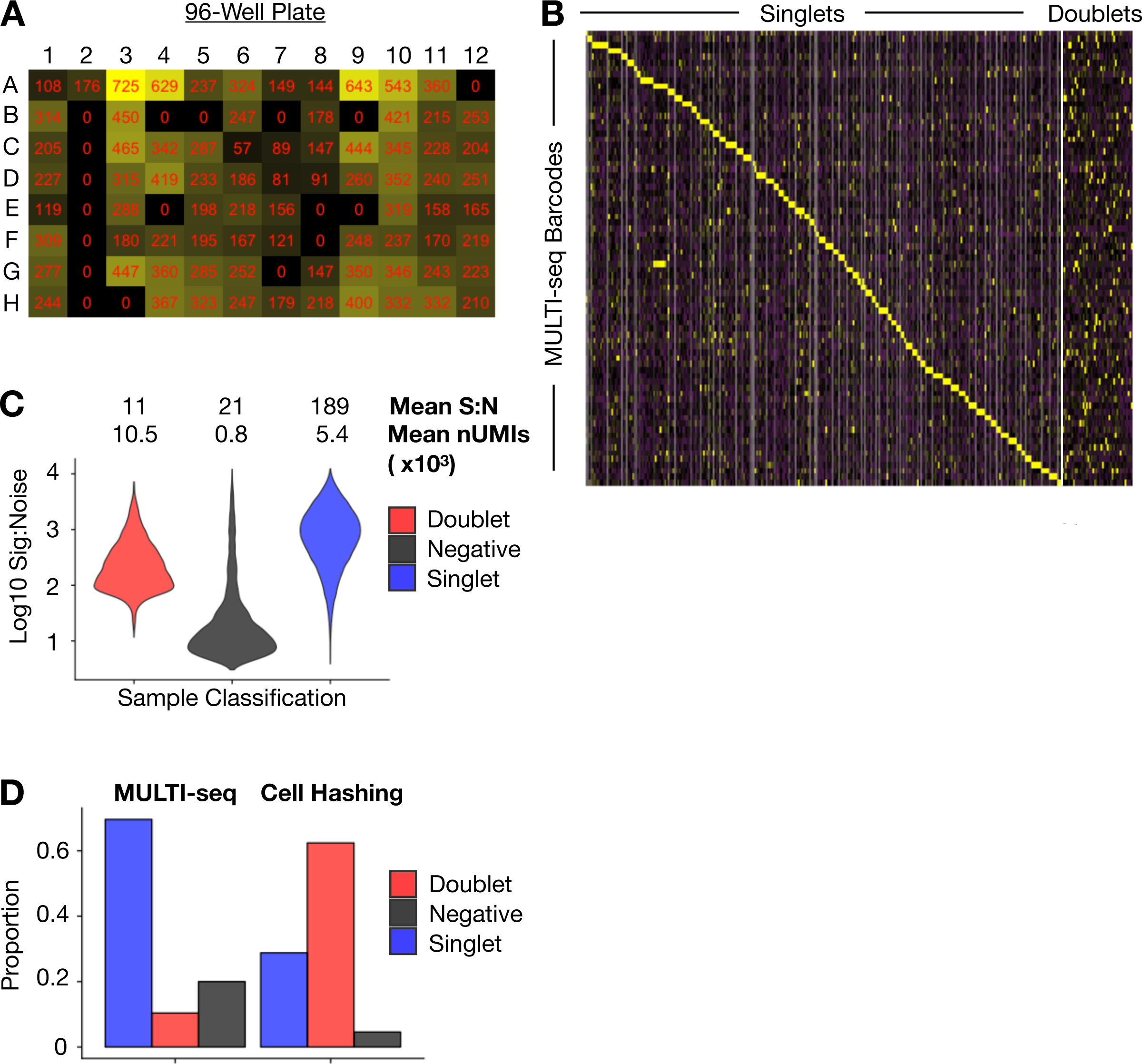
HMEC sample classification results, related to Figure 3. (A) Heatmap showing the number of cells assigned to each sample barcode group arranged according to their position on the 96-well plate utilized during sample preparation. The predominant lack of samples arising from column 2 indicates that technical error during sample preparation likely caused sample drop-outs. (B) Heatmap showing the enrichment within each sample classification group for a single MULTI-seq barcode. Doublets are enriched for multiple barcodes. (C) Violin plots describing the signal:noise for negative cells, doublets and singlets. In singlets, on-target barcodes are an average of 189-fold higher than the most abundant off-target barcode. Doublets have much lower signal:noise but higher total nUMIs, which matches expectations based on the pooling of multiple unique barcodes that occurs during doublet formation. Negative cells exhibit very low total nUMIs, indicating that negative cells were not sufficiently barcoded to enable sample classification. (D) Comparison of sample classification results for the MULTI-seq workflow relative to the Cell Hashing classification strategy (Stoeckius et al., 2017a). Cell Hashing sample classification does not produce as many negative calls, but highly over-estimates the number of doublets.

**Figure S7:**
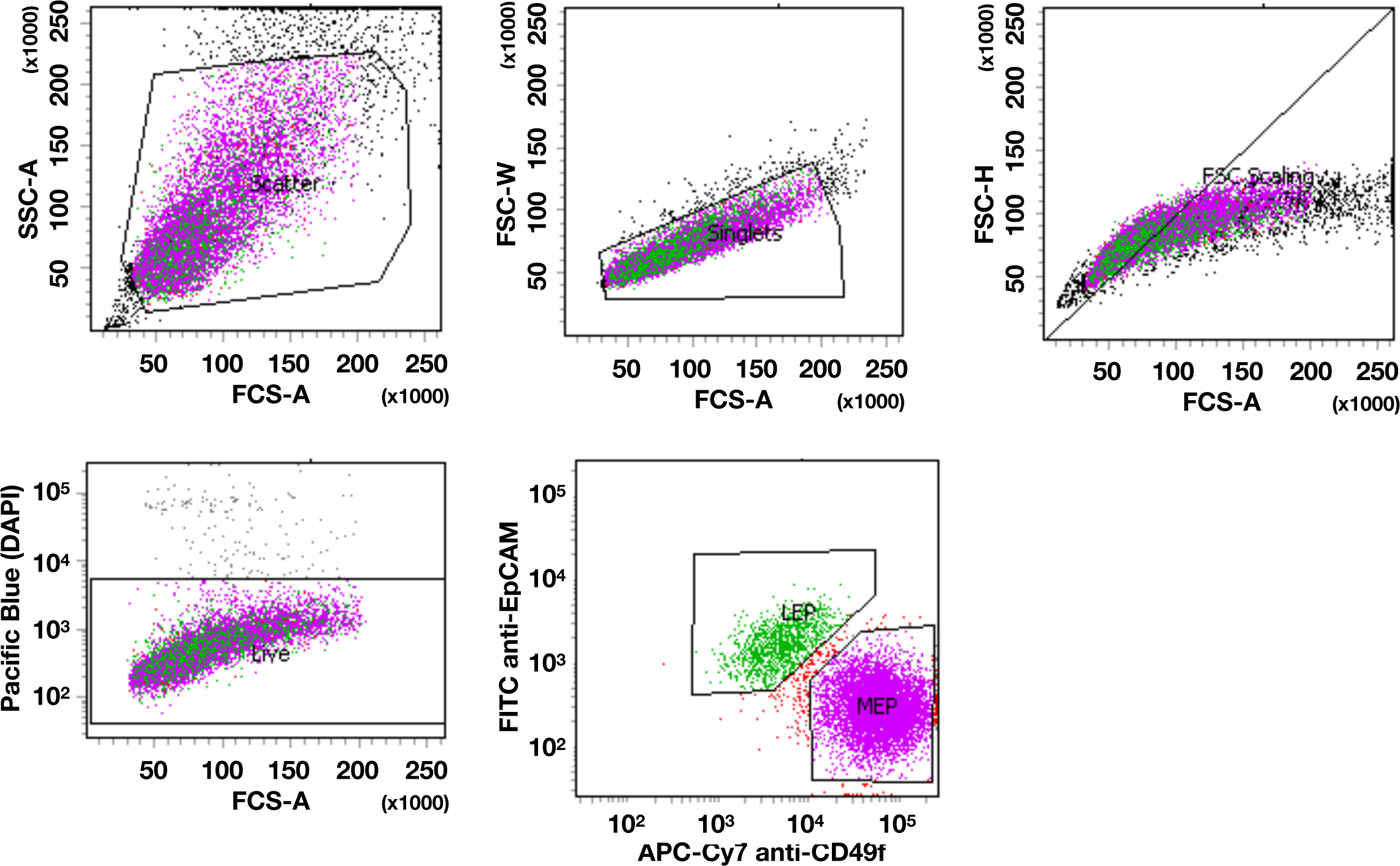
FACS purification of LEP and MEP cells from bulk HMECs, related to Experimental Methods. Bulk HMECs were labeled with FITC anti-EpCAM and APC-Cy7 anti-CD49f to identify and isolate LEPs and MEPs. LEPs are identified as EpCAM high and CD49f low, while MEPs are CD49f high and EpCAM low. Gating strategy causes minor cell type impurities in final sorted population. See methods for full details.

**Figure S8:**
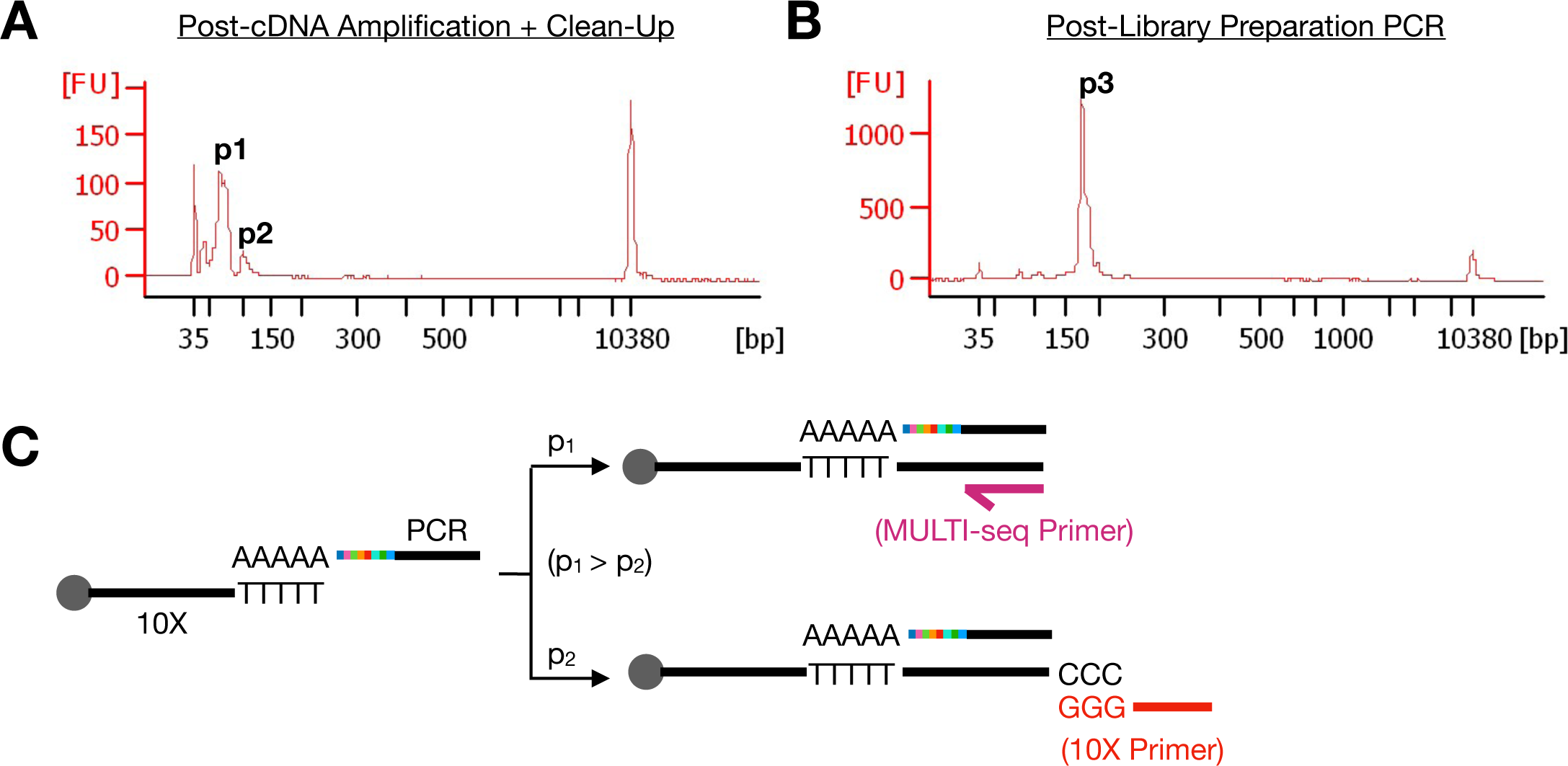
Bioanalyzer traces of representative MULTI-seq barcode library, related to Experimental Methods. (A) Bioanalyzer traces following cDNA amplification and MULTI-seq barcode enrichment using 3.2X SPRI with 1.8X 100% isopropanol exhibits two distinct peaks. The first peak (p1) is an average of 65-70bp in length and likely corresponds to barcodes amplified via the MULTI-seq additive primer. The second peak (p2) is an average of 100bp in length and likely corresponds to barcodes that successfully underwent MMLV-RTase template switching and were subsequently amplified by the standard 10X Genomics Single Cell V2 primer. Considering the low efficiency of template switching relative to processive reverse transcription, the abundance difference of the two peaks fits expectations. (B) Bioanalyzer analysis following library preparation PCR exhibits one distinct peak (p3) with an average length of 173bp, matching expectations. (C) Schematic illustrating the two species of reverse-transcribed MULTI-seq barcodes with and without template switching. Processive reverse-transcription without template switching (p1) is more likely than reverse-transcription with template switching (p2), resulting in relative enrichment of the 65-70bp product following cDNA amplification.

**Table S1:** List of all differentially expressed genes between MULTI-seq barcoded and un-barcoded control cells, related to Figure 1

| GeneID | p (bimod) | Description |
| --- | --- | --- |
| MIF | 0 | Macrophage Migration Inhibitory Factor |
| TOMM5 | 0 | Translocase of outer mitochondrial membrane 5 |
| RPL17 | 0 | Ribosomal Protein L17 |
| NME1-NME2 | 0 | Nucleoside Diphosphate Kinase (NME1-NME2) Read-Through |
| KRTCAP2 | 3.7E-305 | Keratinocyte Associated Protein 2 |
| RPS10 | 1E-277 | Ribosomal subunit protein |
| RPL36A | 2.9E-123 | Ribosomal subunit protein |

**Table S2:** Proportion of mouse and human cells loaded into the 10X microfluidics device relative to in the final dataset, related to Figure 2

| Sample | %Human IN | %Human OUT | %Mouse IN | %Mouse OUT |
| --- | --- | --- | --- | --- |
| A | 66.7 | 91.7 | 33.3 | 8.3 |
| B | 71.4 | 91.9 | 28.6 | 8.1 |
| C | 62.7 | 55.6 | 37.3 | 44.4 |
| D | 17.9 | 4.1 | 81.1 | 95.9 |
| E | 7.6 | 3.5 | 92.4 | 96.5 |
| F | 0.4 | 1.4 | 99.6 | 98.6 |
| G | 0.9 | 2.2 | 99.1 | 97.8 |

**Table S3:**
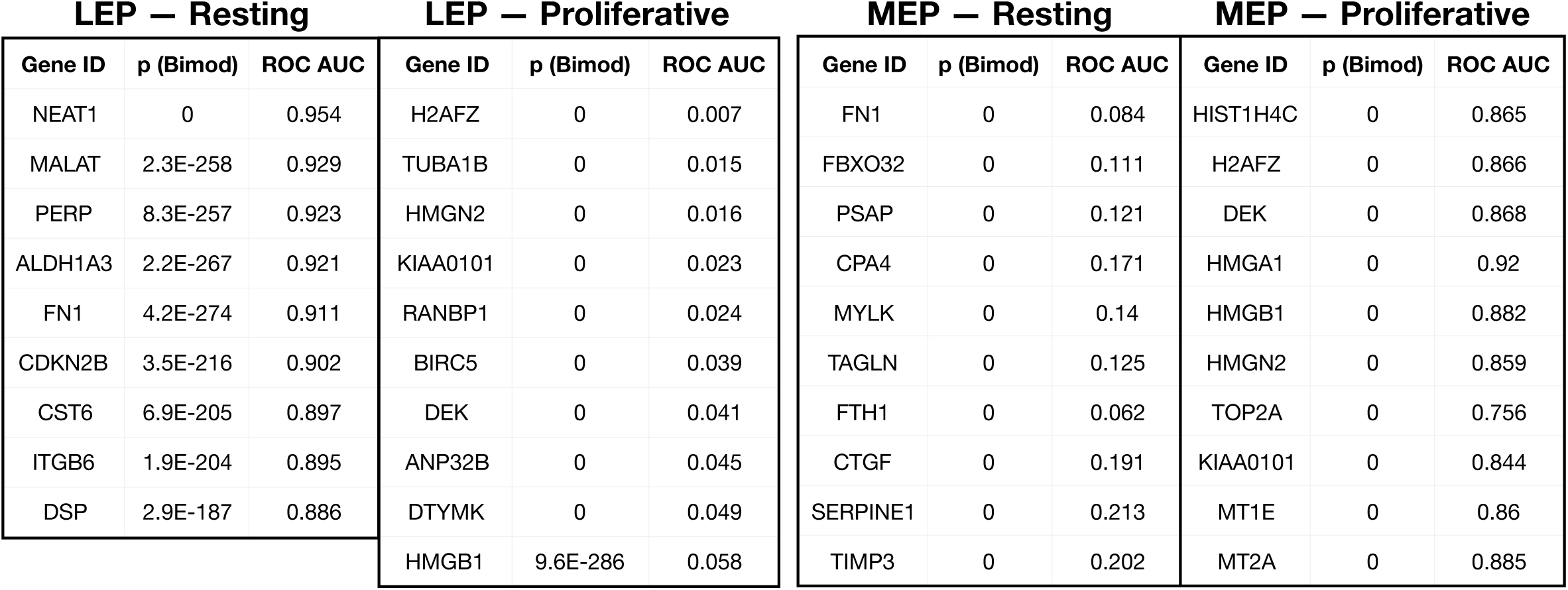
Marker analysis on full MEP and LEP subsets detects proliferative and resting cell states, related to Figure 4

**Table S4:**
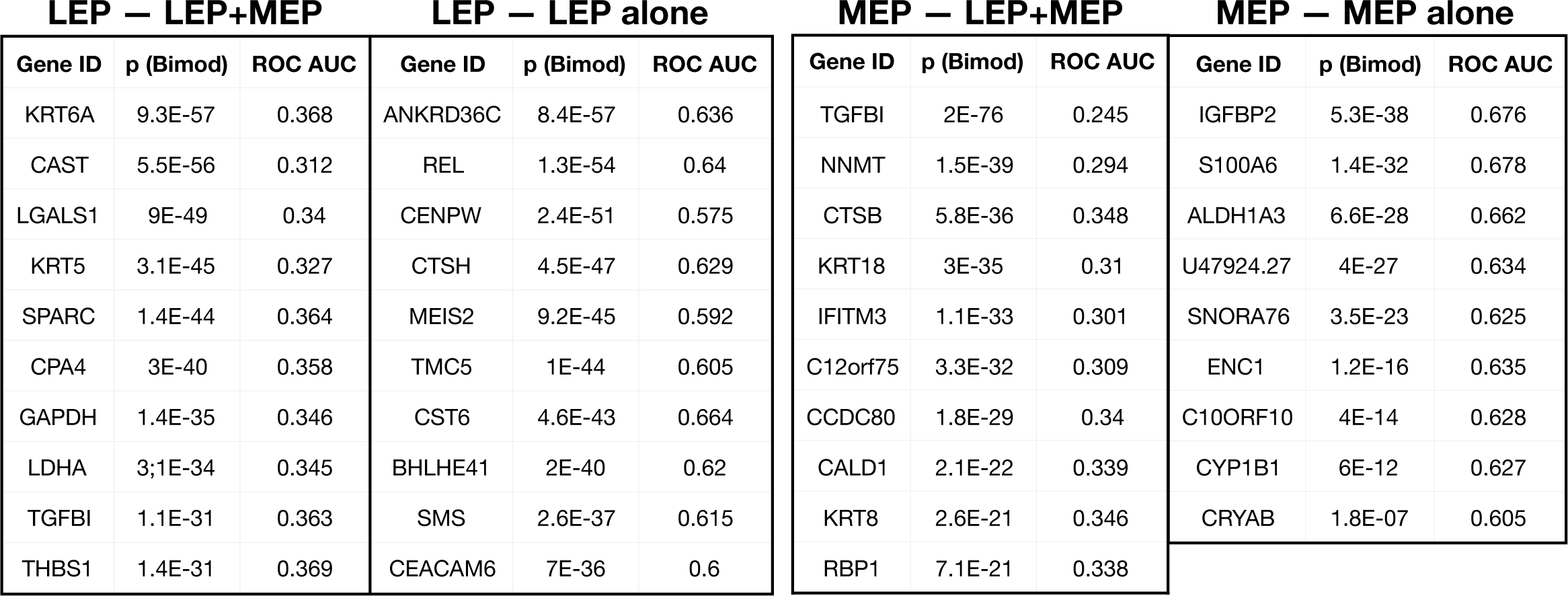
Marker analysis on resting MEPs and LEPs grouped by co-culture status detects TGF-β signaling-associated transcriptional response in co-cultured MEPs and LEPs, related to Figure 4

**Table S5:** Marker analysis on MEPs and resting LEPs grouped by growth factor supplementation detects EGFR-associated transcriptional responses in AREG- and EGF-stimulated cultures, related to Figure 4

| LEP — Control |  |  | LEP — EGFR |  |  | MEP — Control |  |  | MEP — EGFR |  |  |
| --- | --- | --- | --- | --- | --- | --- | --- | --- | --- | --- | --- |
| Gene ID | p (Bimod) | ROC AUC | Gene ID | p (Bimod) | ROC AUC | Gene ID | p (Bimod) | ROC AUC | Gene ID | p (Bimod) | ROC AUC |
| CCDC80 | 2.9E-131 | 0.361 | CCND1 | 2.3E-195 | 0.717 | CALD1 | 0 | 0.296 | ANGTPL4 | 0 | 0.674 |
| FBX032 | 1E-116 | 0.338 | DCBLD2 | 3E-99 | 0.662 | TPM1 | 0 | 0.286 | ADIRF | 0 | 0.698 |
| CENPW | 4.5E-113 | 0.391 | DUSP4 | 1E-92 | 0.660 | HSPB1 | 0 | 0.25 | UPP1 | 0 | 0.706 |
| GPX4 | 4.6E-88 | 0.344 | F3 | 1.1E-69 | 0.635 | MYLK | 0 | 0.361 | SLC20A1 | 0 | 0.701 |
| SPTSSA | 7.8-85 | 0.418 | MALL | 2.5E-69 | 0.636 | TPM2 | 6.9E-305 | 0.319 | FERMT1 | 0 | 0.63 |
| TMC5 | 3.7E-75 | 0.406 | PHLDA1 | 9.3E-66 | 0.636 | TAGLN | 2.7E-278 | 0.324 | HMGA1 | 5.2E-269 | 0.666 |
| MMP7 | 7.1E-72 | 0.382 | THBS1 | 2.5E-59 | 0.626 | MYL9 | 2.2E-277 | 0.327 | PTHLH | 7.9E-195 | 0.636 |
| REL | 4.7E-71 | 0.413 | LAMC2 | 3.7E-44 | 0.602 | KRT17 | 1E-269 | 0.332 | TFPI2 | 3.3E-180 | 0.596 |
| RARRES1 | 2.3E-61 | 0.368 | KRT81 | 7.8E-38 | 0.605 | KLK7 | 1.97E-251 | 0.353 | S100A6 | 3.9E-177 | 0.63 |
| NEAT1 | 2E-60 | 0.370 | SERPINE1 | 2.1E-16 | 0.565 | CDC42EP3 | 1.6-242 | 0.382 | G032 | 2.6E-145 | 0.59 |

